# Development of a novel high-throughput screen for the identification of new inhibitors of protein S-acylation

**DOI:** 10.1101/2022.03.17.484726

**Authors:** Christine Salaun, Hiroya Takizawa, Alex Galindo, Kevin R. Munro, Jayde McLellan, Isamu Sugimoto, Tomotaka Okino, Nicholas C.O. Tomkinson, Luke H. Chamberlain

## Abstract

Protein S-acylation is a reversible post-translational modification that modulates the localisation and function of many cellular proteins. S-acylation is mediated by a family of zinc finger DHHC domain-containing proteins encoded by 23 distinct *ZDHHC* genes in the human genome. These enzymes catalyse S-acylation in a two-step process involving “auto-acylation” of the cysteine residue in the catalytic DHHC motif followed by transfer of the acyl chain to a substrate cysteine. S-acylation is essential for many fundamental physiological processes, and there is growing interest in zDHHC enzymes as novel drug targets for a range of disorders. However, there is currently a lack of chemical modulators of S-acylation either for use as tool compounds or for potential development for therapeutic purposes. In this study, we developed and implemented a novel FRET-based high throughput assay for the discovery of compounds that interfere with auto-acylation of zDHHC2, an enzyme that is implicated in neuronal S-acylation pathways. A screen of >350,000 compounds identified two related tetrazole containing compounds (TTZ-1 and -2) that inhibited both zDHHC2 auto-acylation and substrate S-acylation in cell-free systems. Furthermore, these compounds were also active in HEK293T cells, where they inhibited substrate S-acylation mediated by different zDHHC enzymes, with some apparent isoform selectivity. Resynthesis of the hit compounds confirmed their activity, providing sufficient quantities of material for further investigations. The assays developed herein provide novel strategies to screen for zDHHC inhibitors, and the identified compounds add to the chemical toolbox for interrogating the cellular activities of S-acylation and zDHHC enzymes.

## INTRODUCTION

S-acylation (or S-palmitoylation) is the post-translational attachment of long chain fatty acids (predominantly palmitates) to cysteines *via* a labile thioester bond (1–4). S-acylation is reversible, making it a dynamic post-translational modification that is able to regulate the stability, localisation, activity and function of a broad range of proteins. Around 10% of the human proteome might be modified by S-acylation (5), and this reaction is catalysed by a family of twenty-three zinc finger DHHC domain-containing (zDHHC) enzymes, defined by a 51-amino acid cysteine-rich domain (CRD) with an Asp-His-His-Cys (DHHC) motif (6–8). zDHHC enzymes use a two-step ping-pong mechanism to mediate S-acylation (9–11). The first step is the auto-acylation of the enzyme that involves the attack on the acyl-CoA by the catalytic cysteine within the DHHC sequence. The second step is the transfer of the acyl chain from the enzyme to a cysteine residue of the target substrate. The crystal structures of zDHHC20 and zDHHC15 have been reported recently (11,12), showing that the four transmembrane domains of these enzymes arrange into a tepee-like structure, providing a hydrophobic cavity for the fatty acyl chain that becomes attached to the DHHC motif cysteine during the auto-acylation step of the enzyme reaction. A preference of some zDHHC enzymes for specific fatty acid substrates was revealed by *in vitro* (9) and *in cellulo* studies (13) before being confirmed following the resolution of the crystal structure of zDHHC20 bound to a fatty acid chain (11,12). The amino-acid residues lining the hydrophobic cavity indeed act as a molecular ruler for the length of the fatty acid chain that can be accommodated. In addition to the catalytic DHHC domain, conserved PaCCT (palmitoyltransferase conserved C-terminus) and TTxE (Thr-Thr-X-Glu) motifs in the C-terminus of zDHHC20 were suggested to be important for enzyme function by forming key contacts with the catalytic site and other parts of the protein (8,11,14).

Analysis of the curated S-acylated proteome indicates that 41% of synaptic genes encode proteins that undergo S-acylation (15). Broad changes in S-acylation levels are induced by modulation of neuronal activity, highlighting a fundamental role of S-acylation in regulating synaptic functions (16). Alterations in the balance between neuronal excitation and inhibition are associated with several neuropsychiatric disorders such as autism, epilepsy and schizophrenia (17). Given the large number of neuronal substrates and the dynamic nature of S-acylation it is not surprising that this modification has been linked with various neurological disorders (18–21). Genome-wide association analyses have identified *ZDHHC2* as a candidate gene for schizophrenia (22,23) as well as Parkinson’s disease (24). A targeted study identified rare mutations within the *ZDHHC2* coding sequence of Han Chinese patients suffering from schizophrenia (23). It was also suggested that the presence of zDHHC2 protects dopaminergic neurons from death during the course of Parkinson’s disease (25). There is strong evidence linking mutations in *ZDHHC9* (26,27) with intellectual disability. A recent study revealed an elevated expression of zDHHC8 in the brain tissue of patients suffering from temporal lobe epilepsy (28), and there is also a possible (although still controversial) association between *ZDHHC8* polymorphism and schizophrenia (19,21). S-acylation deficits have been suggested in other neurological disorders (such as Alzheimer’s or Huntington’s disease) although demonstrated contributions of zDHHC enzymes to these are still missing (19,21). In addition, to neurological disorders, there is also evidence linking zDHHC enzymes to diabetes and cancer (2).

A major obstacle to progress of the S-acylation field is a lack of selective chemical inhibitors or modulators of zDHHC enzymes (29–33). Three main lipid-based chemical compounds have long been used for their ability to inhibit S-acylation (Supp Fig 1). The most widely used compound is 2-bromopalmitic acid (2BP), whereas cerulenin and tunicamycin have been used less frequently. However, none of these compounds is thought to be selective for a specific zDHHC enzyme(s) and many of the effects of these compounds may be entirely non-specific. Tunicamycin is primarily known as an inhibitor of N-linked glycosylation, and cerulenin has been shown to inhibit various fatty acid synthases (34). Although 2BP has been shown to irreversibly block the activity of purified zDHHC enzymes (6,35) and bind to zDHHC enzymes in cells (36,37), it has also been found to be highly promiscuous (38), labelling hundreds of cellular proteins (36,37) and therefore has potential pleiotropic effects on cell metabolism (29). An effort to develop a broad spectrum inhibitor of zDHHC enzymes that would overcome the major drawbacks of 2BP led to the synthesis of CMA (*N*-cyanomethyl-*N*-myracrylamide) which utilises an acrylamide warhead linked to the same fatty-acid chain as 2BP (33,39). This promising new compound has been shown to inhibit the activity of DHHC20 *in vitro* (IC_50_ of 1.35 +/- 0.26 μM) and of many DHHCs *in cellulo* with a low level of toxicity.

Several screens were developed in an attempt to find more specific inhibitors of zDHHC-mediated S-acylation (31,33,40,41). Five (non-lipid based) compounds were identified from the DIVERSet collection (ChemBridge Corporation, San Diego, CA) for their ability to reduce peptide substrate S-acylation (40). However, further investigation showed that the potency and selectivity of four of these compounds were not recapitulated with purified enzymes and their cognate substrates. Only one of the compounds (Compound V) directly (and reversibly) inhibited the acyltransferase activity of four different yeast and human zDHHC enzymes, and may therefore be a general inhibitor of zDHHC enzymes (35). Inhibitors of the zDHHC enzyme Erf2 (the yeast ortholog of the mammalian zDHHC9 enzyme) were identified from a large scaffold-ranking library (41). The hit compounds against Erf2 were based on a bis-cyclic piperazine backbone. Interestingly, the identified compounds were recently shown to inhibit zDHHC9 mediated S-acylation of the spike protein of the Severe Acute Respiratory Syndrome Coronavirus 2 (SARS-CoV-2), the virus responsible for the Coronavirus disease 2019 pandemic, and to reduce virus infectivity (42). Protein palmitoylation inhibition has indeed been identified as a therapeutic target against SARS-CoV-2 (43). Finally, Qiu *et al* recently reported the serendipitous discovery of Artemisinin (an antimalarial sesquiterpene lactone) as a zDHHC6 covalent inhibitor that attenuates NRas (and other zDHHC6 substrates) S-acylation without altering global S-acylation (44), suggesting that the selective inhibition of S-acylation enzymes might be achievable.

To advance efforts to identify novel S-acylation inhibitors, we have developed a high-throughput FRET-based autoacylation assay to screen >350,000 chemicals for activity against zDHHC2 autoacylation. Furthermore, selected hits were further assessed using novel *in vitro* and established *in cellulo* click chemistry-based substrate S-acylation assays. Two related compounds were identified that displayed activity against zDHHC2 in all three of these different assay formats. The compounds also displayed partial enzyme isoform selectivity in cell-based assays. This study thus expands the available repertoire of assay platforms and chemical inhibitors to advance S-acylation research.

## RESULTS

### Development of a novel FRET-based high-throughput screen to identify chemical modulators of zDHHC2 autoacylation

S-acylation is a two-step process involving enzyme “auto-acylation” (at the cysteine of the DHHC motif) followed by transfer of the acyl chain to a substrate protein. To identify potential inhibitors of S-acylation, we developed an assay measuring enzyme auto-acylation and focused on zDHHC2 given its prominent role in synaptic function in the nervous system (Figure 1A). The assay monitors time-resolved fluorescence resonance energy transfer (TR-FRET) between the energy donor terbium crypate (attached to an antibody recognising HA-tagged zDHHC2) and the acceptor fluorophore nitrobenzoxadiazole NBD (attached to the lipid group). The excitation of the energy donor terbium cryptate (360 nm) followed by its emission at 480 nm results in energy transfer to the acceptor NBD and emission at 539 nm. Enzyme auto-acylation by NBD-conjugated palmitoyl-CoA increases the proximity of the terbium cryptate and NBD, leading to an increase in FRET signal. The resulting interaction is quantified as the ratiometric measurement of NBD (539 nm) over terbium (480 nm) emission and constitutes the TR-FRET signal.

**Figure 1.**
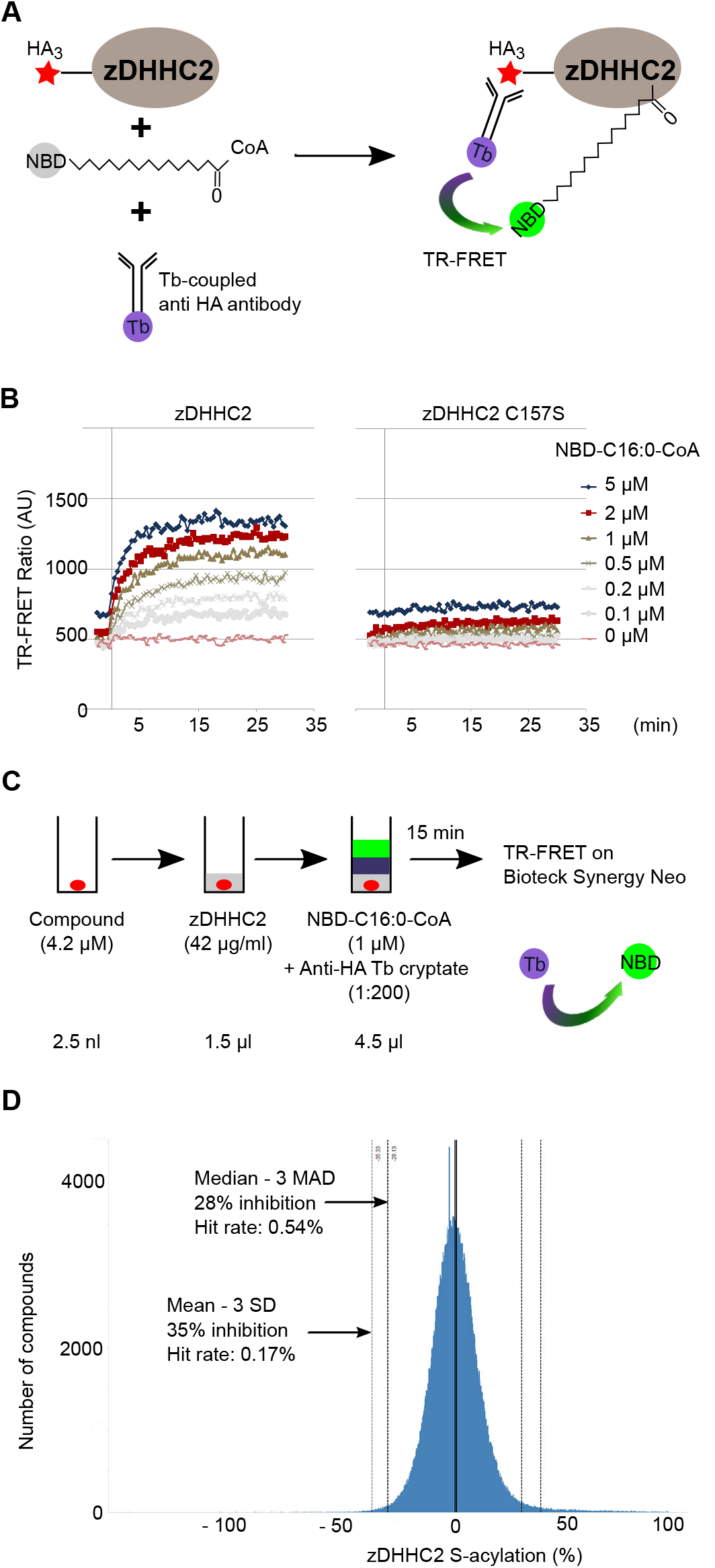
High throughput screen (HTS) for the discovery of chemical compounds modulating human zDHHC2 autoacylation activity: TR-FRET-based assay. **A.** Experimental design of the HTS. HA-tagged zDHHC2 enzyme expressed at the surface of lipoparticles was incubated for 15 minutes at room temperature (RT) with NBD-tagged palmitoyl-CoA and an anti-HA antibody coupled to terbium cryptate (Tb). Autoacylation of zDHHC2 on its catalytic cysteine brings closer the FRET (Fluorescence Resonance Energy Transfer) donor Terbium crypate bound to the anti-HA antibody to the NBD fluorophore coupled to the palmitate and triggers FRET that can be followed over time (Time Resolved FRET, TR-FRET). **B.** TR-FRET was measured with increasing concentrations of NBD-palmitoyl-CoA (NBD-C16:0-CoA) and either WT zDHHC2 (*left panel*) or catalytically dead zDHHC2 C157S (*right panel*) over a period of 35 minutes. **C.** Experimental conditions of the HTS screen. 350,000 compounds were incubated individually in 1536 well plates at a concentration of 4.2 μM with zDHHC2 lipoparticles (42 μg/ml stock) for a few minutes before the addition of a mix containing the Terbium (Tb) cryptate-tagged anti-HA antibody (1:200) and 1 μM of NBD-palmitoyl-CoA (NBD-C16:0-CoA). Time-resolved FRET was measured after a 15 minute incubation at room temperature on a Biotek Synergy Neo apparatus. Results are given as a ratio between the FRET signal measured over a background baseline. **D.** Histogram representation of ONO library screen showing the percentage of activity of zDHHC2 (as measured by TR-FRET) in the presence of each compound. The graph represents the number of compounds that are either activating (0 to 100) or inhibiting (0 to -100) zDHHC2 activity. Compounds showing over 35% inhibition (Mean – 3 Standard Deviation SD) were considered as hits.

Human zDHHC2 enzyme for use in this assay was produced with the Expi293 MembranePro™ expression system (Invitrogen), which enables membrane proteins to be displayed in a native context on highly enriched lipoparticles that are released in the supernatant of the transfected mammalian cells.

An example of the TR-FRET measured using this assay system is shown in Figure 1B. Auto-acylation of zDHHC2, detected by TR-FRET signal, was shown to increase with increasing concentration of NBD-palmitoyl-CoA. Importantly, the TR-FRET signal was substantially reduced when a catalytically-dead form of zDHHC2 with a substitution of the cysteine in the DHHC motif (C157S) was used, demonstrating that the FRET signal faithfully reports the auto-acylation of the enzyme.

Following optimisation, the assay was miniaturized to a 1536-well plate format. A pilot screen was then performed at two concentrations (12.5 and 4.2 μM) using ONO pilot library containing a total of 15,792 compounds. RZ’ score was calculated to be between 0.72 and 0.80 for each plate showing that the assay design was robust. Figure 1C outlines the final conditions chosen for the screen, which were then used to screen a HTS library of >350,000 compounds at a concentration of 4.2 μM (Figure 1D).

One hundred and fifty compounds that displayed activity in the TR-FRET assay screen were selected for further evaluation. Specifically, their activity was confirmed using freshly-dissolved powders and at two different concentrations (12.5 and 4.2 μM). The results presented focus on two of these compounds (Tetrazole (TTZ) -1 and -2), which were subsequently shown to have an activity on cells. The chemical structure of these two compounds is shown in Figure 2. Table 1 presents data obtained from the HTS and confirmation screens. zDHHC2 auto-acylation was inhibited by 61% by TTZ-1 and 69% by TTZ-2 at a concentration of 12.5 μM.

**Figure 2.**
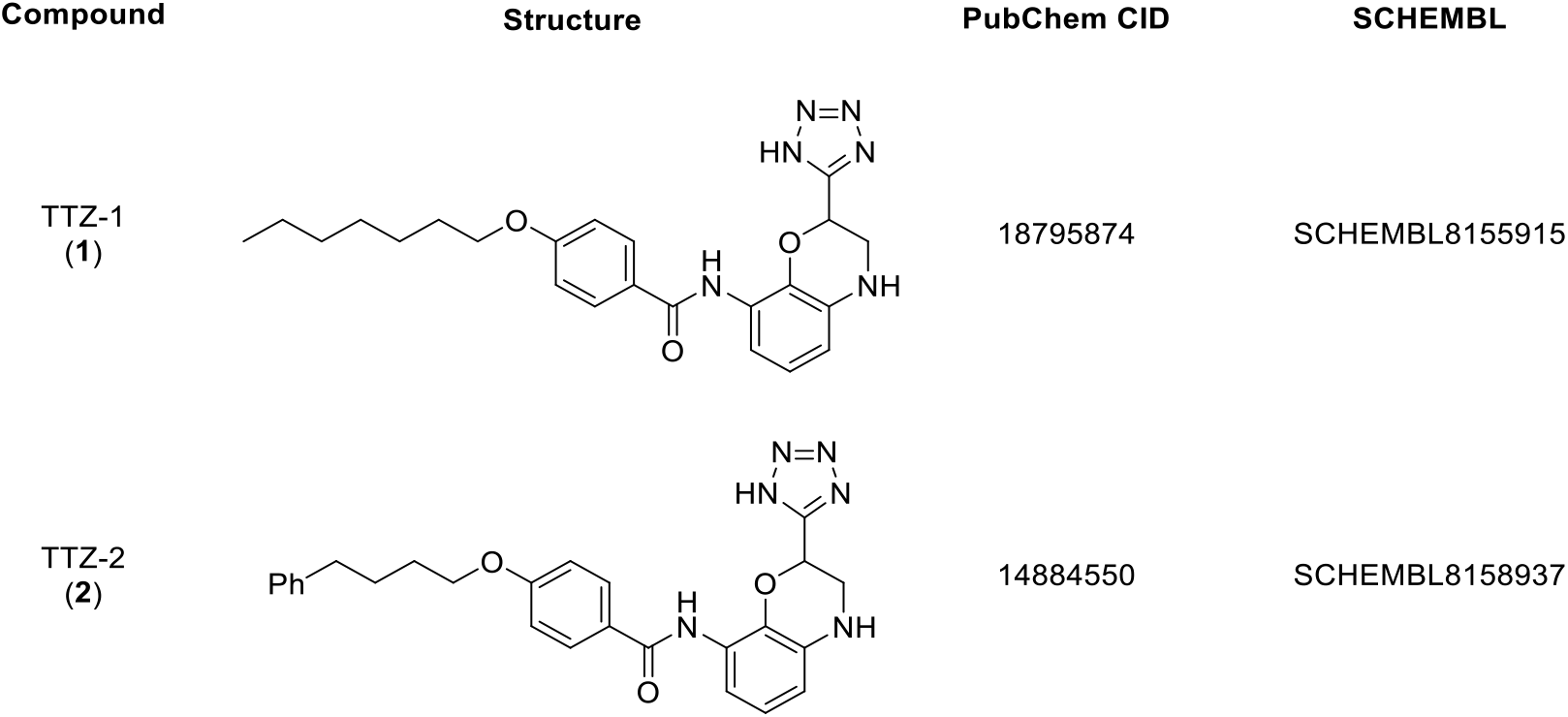
Chemical structure of TTZ-1 and TTZ-2.

**Table 1.**
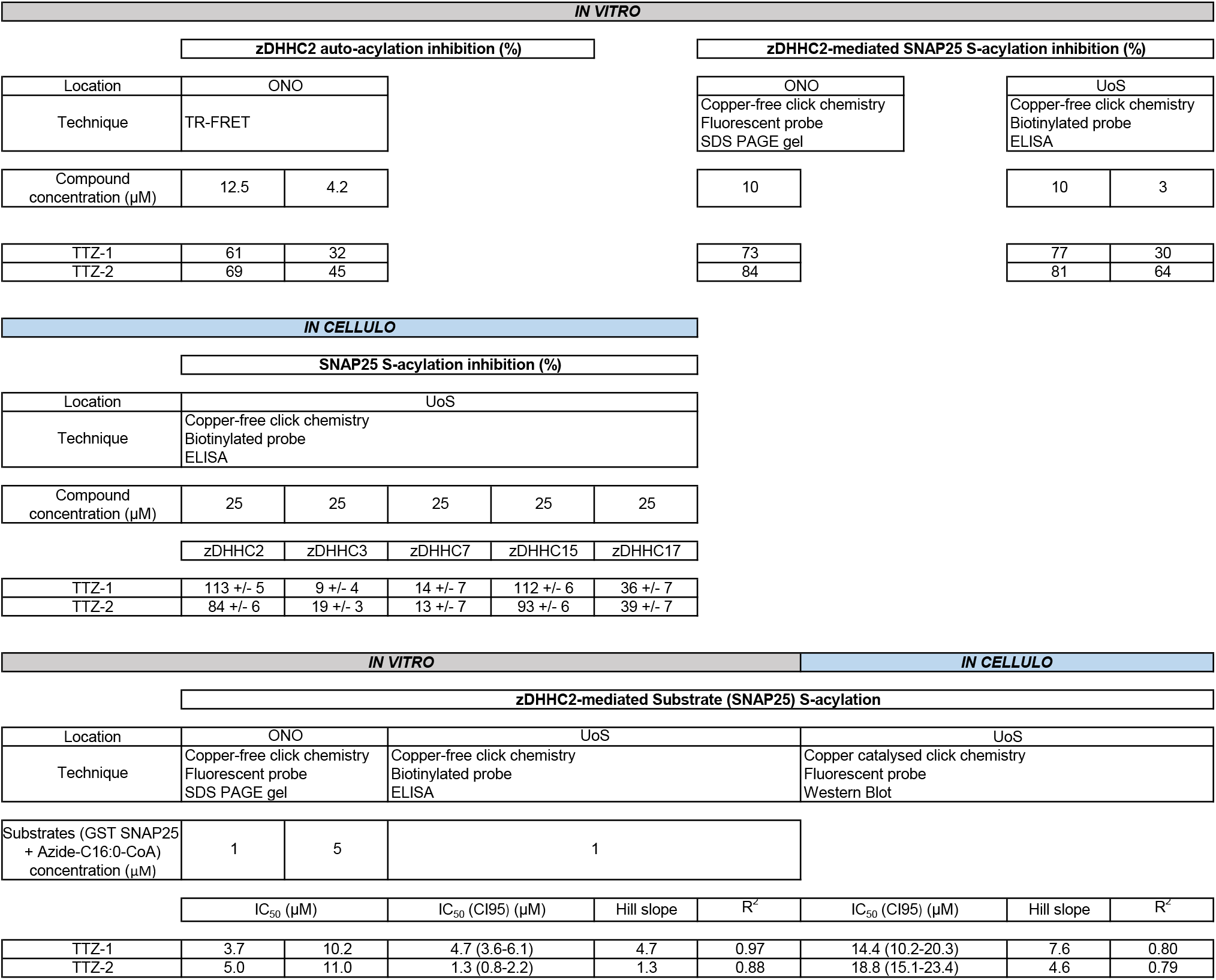
Summary of data obtained at both research sites for compounds TTZ-1 and TTZ-2. The table gives the location of the research (either Ono or University of Strathclyde UoS), the technique used and the numerical data generated, both for *in vitro* and in HEK293T cells (*in cellulo*). The upper 3 tables summarize the efficiency of inhibition achieved (in %) for the auto-acylation of the enzyme(s) and for the substrate S-acylation, at various concentrations of compounds. The lower 2 tables gather data from IC_50_ determination.

### Development of a novel ELISA assay to detect in vitro substrate S-acylation by purified human zDHHC2

We next assessed the ability of the selected compounds to inhibit zDHHC2-mediated substrate S-acylation *in vitro*. For this, we designed an ELISA assay in a 96-well plate format. His_6_-tagged human zDHHC2 enzyme was purified from insect cells on Ni^2+^-NTA affinity resin according to methods developed in (9) (Figure 3A). GST-tagged SNAP25, a reported substrate of zDHHC2 (45) was then incubated with the purified zDHHC2 enzyme and azide-C16:0-CoA for 30 to 45 min (GST was used as a control). After treatment with 50 mM N-ethyl maleimide (NEM) for 2 hours to block free cysteines, the azide tagged palmitate that had been incorporated into GST-SNAP25 and zDHHC2 was reacted with DBCO-PEG4-Biotin in a copper-free cycloaddition reaction. Samples were then analysed by immunoblotting with NeutrAvidin-800 to detect biotin (and hence S-acylation) and an anti-GST antibody (IR680) to detect GST/GST-SNAP25. Figure 3B shows that this assay detects the S-acylation of GST-SNAP25 (band above 47 kDa) but not GST. The NeutrAvidin-800 signal also revealed the auto-acylated enzyme (band above 36 kDa).

**Figure 3.**
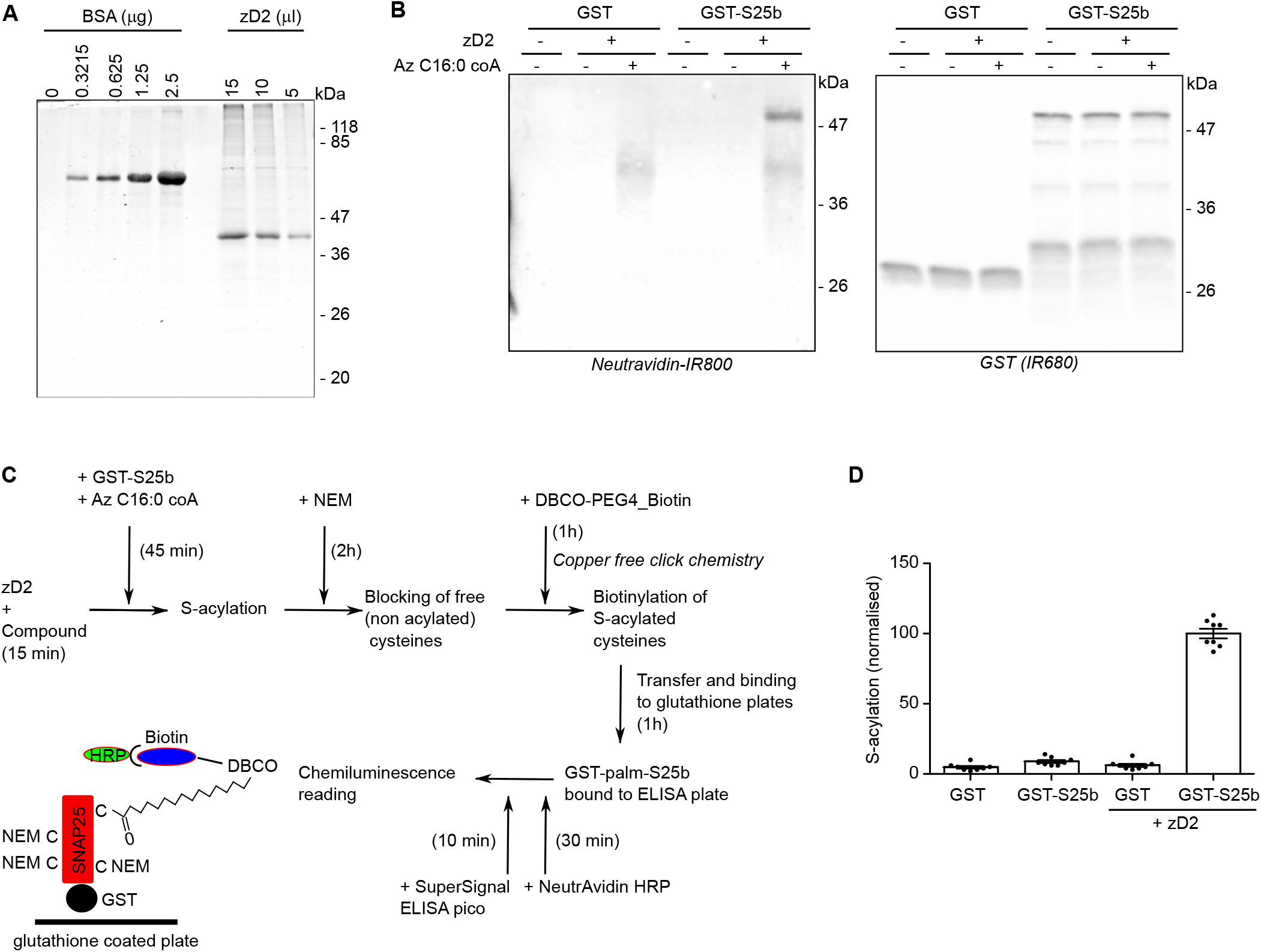
Purification of active human zDHHC2 enzyme from insect cells and development of an ELISA assay for the detection of substrate S-acylation. **A.** Purification of human zDHHC2. His-tagged human zDHHC2 (zD2) was expressed with a baculovirus system in Sf9 TriEX insect cells and purified using Ni^2+-^NTA resin. Aliquots were resolved on a SDS PAGE gel together with increasing amounts of Bovine Serum Albumin (BSA) standards. Proteins were revealed with a staining agent (Instant Blue). The main band migrating between 36 and 47 kDa is at the estimated size for tagged zDHHC2. Position of the molecular mass marker (in kDa) are indicated on the right side of the gel. **B.***In vitro* S-acylation with purified zDHHC2 and detection by copper-free click chemistry. The zDHHC2 substrate GST-SNAP25b (GST-S25b) or a control GST protein (both at 1 μM final) were incubated either with (+) or without (-) zDHHC2 (zD2) (0.075 μM final), and either with (+) or without (-) 1 μM Azide-C16:0-CoA (Az-C16:0-CoA) in acylation buffer for 45 minutes at 25°C. Free (non-acylated cysteines) were then blocked for 2 hours with 50 mM NEM. Copper-free click chemistry was then started by the addition of 5 μM DBCO-PEG4-Biotin in PBS. Samples were incubated for 1 hour at RT and diluted 5 times by the addition of PBS before being resolved on a SDS PAGE gel and transferred to a nitrocellulose membrane. Biotinylated proteins were revealed by incubating the membrane with NeutrAvidin-IR800 whereas the GST-tagged proteins were detected with an anti GST antibody (IR680). Position of the molecular mass marker (in kDa) are indicated on the right side of the membranes. **C.** Schematics of the ELISA assay protocol. **D.** Samples were treated as in B and analysed with the S-acylation ELISA assay as schematised in C. The graph represents the mean +/- SEM of the normalized ELISA values. Each circle represents one value; values were gathered from 4 independent *in vitro* S-acylation experiments performed in duplicate.

This assay was further developed into an ELISA format to more precisely quantify SNAP25 S-acylation (Figure 3C). *In vitro* S-acylation and click chemistry labelling were carried out in uncoated 96 well plates. The processed samples were then diluted with PBS and bound for 1 h to glutathione plates before being incubated with NeutrAvidin-peroxidase (HRP). Chemiluminescence signal was then measured following a 10 min incubation with a peroxidase substrate. Compounds tested in this ELISA assay were pre-incubated with the purified zDHHC2 enzyme for 15 min at room temperature prior to the addition of substrates. Figure 3D shows the normalised chemiluminescent signal detected with the ELISA assay. Background levels of signal were detected with GST or GST-SNAP25 alone (with no added zDHHC2) (4.8 +/- 0.9 and 8.9 +/- 1.0, respectively), or with GST + zDHHC2 (6.1 +/-1.1), whereas there was a strong signal for samples incubated with GST-SNAP25 and the purified enzyme (100 +/- 3.4). The ELISA assay was therefore suitable for testing the activity of the identified compounds on the S-acylation of SNAP-25 by purified zDHHC2.

### TTZ-1 and TTZ-2 inhibit zDHHC2-mediated SNAP25 S-acylation in vitro with IC_50_ values below 5 μM

The ability of TTZ-1 and -2 to inhibit SNAP25 S-acylation by purified zDHHC2 *in vitro* was then determined by calculating IC_50_ values (with concentrations ranging from 0.3 to 30 μM). Figure 4 shows that both compounds inhibited S-acylation of SNAP25 by zDHHC2 with an IC_50_ below 5 μM (Table 1). Selected samples were also analysed by SDS-PAGE and western blotting, which confirmed the inhibitory activity of both compounds (at 3 or 10 μM) on SNAP25 S-acylation (Figure 4C). Importantly, this confirms that the decreased signal detected with these compounds in the ELISA assay reliably reports on reduced S-acylation rather than any non-specific effects.

**Figure 4.**
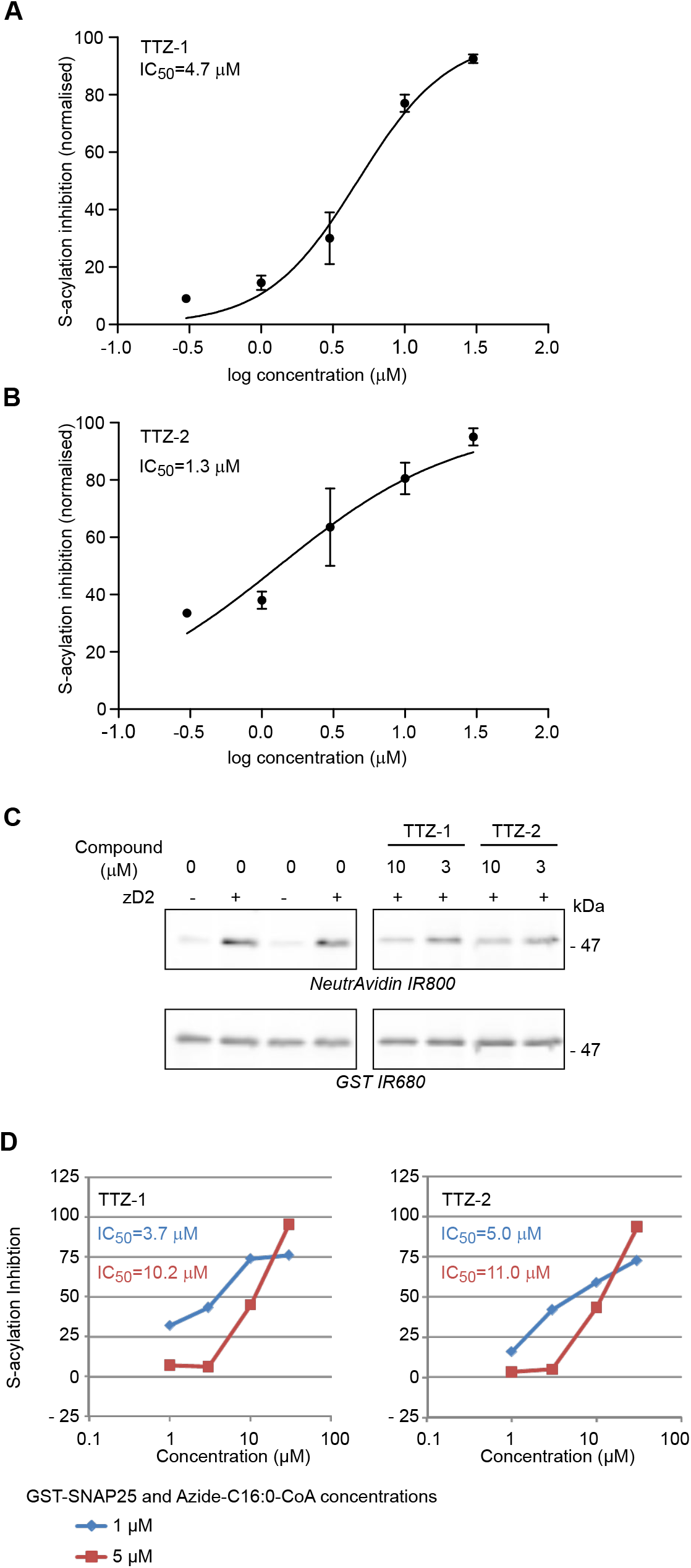
Activity of TTZ-1 and TTZ-2 against SNAP25 S-acylation by purified human zDHHC2 *in vitro*. Increasing concentrations of TTZ-1 (**A**) and TTZ-2 (**B**) compounds or DMSO (as a control) were preincubated with purified zDHHC2 (zD2) for 15 min at room temperature. Azide-C16:0-CoA and GST-SNAP25b were then added and S-acylation was measured as described in figure 3C. The mean ELISA value of a control sample (GST-SNAP25b without zDHHC2; basal background values of the ELISA assay) was subtracted from all the values which were then normalised to the mean value of the control sample GST-SNAP25b + zDHHC2 treated with DMSO. Data shown on the graphs are mean +/- SEM of the percentage of inhibition of zDHHC2 mediated GST-SNAP25b S-acylation for 2 independent experiments. Data were analysed and fit to a nonlinear curve using the log(inhibitor) versus normalised response (variable slope) equation (GraphPad). **C.** Selected samples from the S-acylation reactions from A and B were resolved on a SDS PAGE gel, transferred to a nitrocellulose membrane and probed with NeutrAvidin 800 and an anti GST antibody (IR 680). Molecular mass markers are indicated on the right side of the blots. The immunoblots shown are from the same membrane but have lanes removed that contain samples that are not relevant to this study. **D**. S-acylation assays performed at Ono’s research location with insect cell purified zDHHC2, either 1 μM (Blue line on graphs) or 5 μM (Red line) of both GST-SNAP25 and Azide-C16:0-CoA and increasing concentrations of either TTZ-1 (*left panel*) or -2 (*right panel*). S-acylation was detected by copper-free click chemistry with a fluorescent dye (DBCO-PEG_4_-Fluor 545); samples were resolved on a SDS PAGE gel and fluorescence was quantified within the gel. Results are presented as in A and B.

The inhibitory effects of TTZ-1 and -2 on SNAP-25 S-acylation was also verified using a slightly modified assay. Here, *in vitro* S-acylation was carried out as previously described but in this case S-acylation efficiency was detected by copper-free cycloaddition of a fluorescent DBCO probe. Samples were run on a SDS gel and the fluorescence signal was quantified within the gel. The efficiency of inhibition obtained for compounds used at 10 μM were very similar using these assays (Table 1). Additionally, performing the assay using a higher concentration of substrates resulted in a higher IC_50_ value for both TTZ-1 and TTZ-2 suggesting that these compounds may be competitive inhibitors of zDHHC2 (Figure 4D and Table 1).

### TTZ-1 and -2 inhibit zDHHC2 mediated SNAP-25 S-acylation in cells with IC_50_ values below 20 μM

The inhibitory activity of TTZ-1 and -2 on cells expressing SNAP25 and zDHHC2 was also examined in intact cells. HEK293T cells were transfected with plasmids encoding GFP-tagged SNAP25b and HA-tagged zDHHC2. Cells were then pre-incubated for 4 h with either DMSO as a vehicle control, or increasing concentrations of TTZ-1/2 (diluted in serum-free medium) before being metabolically labelled for a further 4 h with C16:0-azide. Samples were then lysed and processed by copper-catalysed click chemistry to reveal the extent of S-acylation. Figures 5A and 5B show that both compounds inhibited SNAP25 S-acylation catalysed by zDHHC2, with IC_50_ values of approximately15 μM for TTZ-1 and 19 μM for TTZ-2.

**Figure 5.**
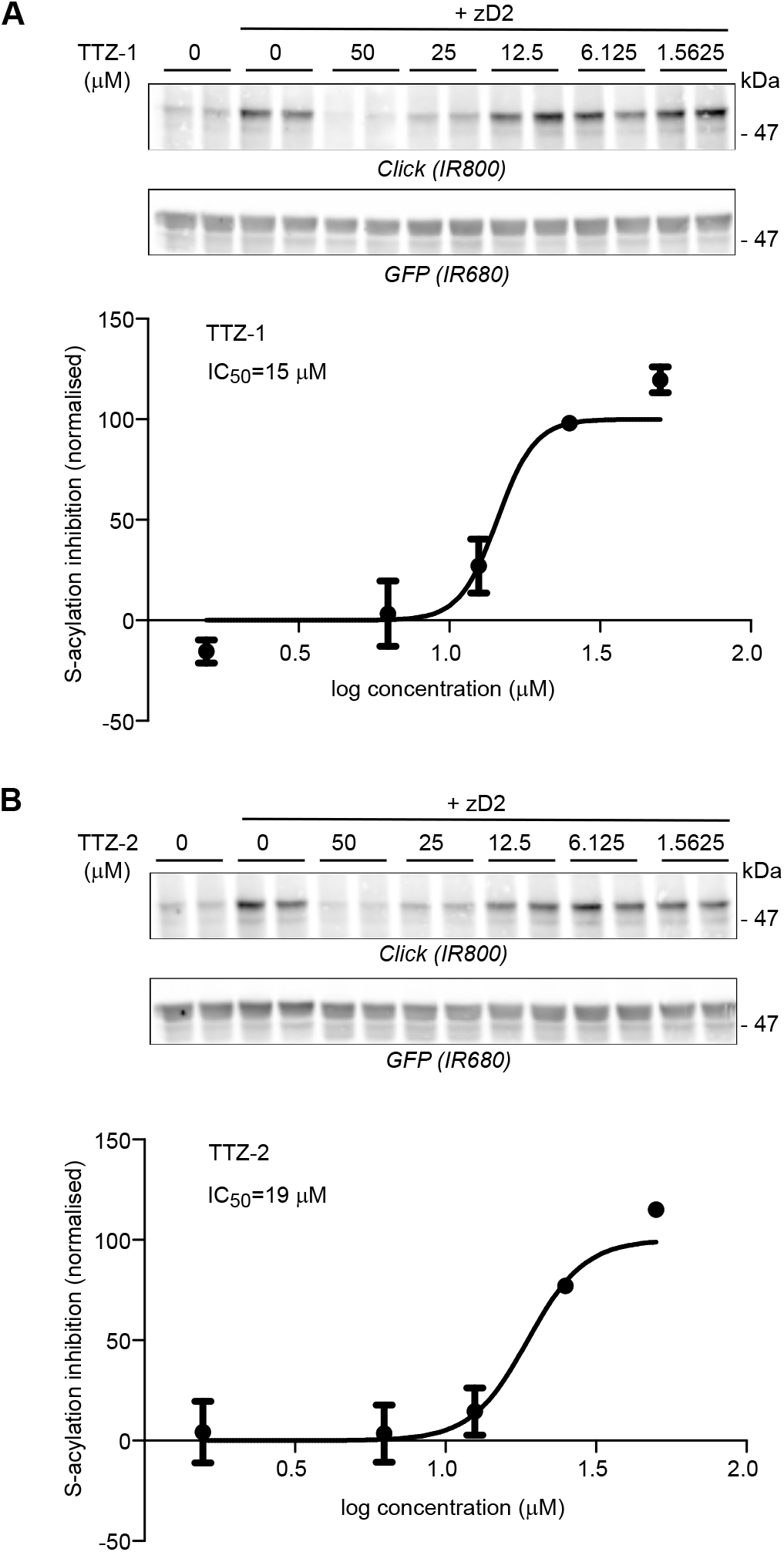
Activity of TTZ-1 and TTZ-2 against SNAP25 S-acylation by human zDHHC2 expressed in mammalian cells. HEK293T cells were transfected with a plasmid encoding GFP-SNAP25b together with a plasmid encoding human zDHHC2 (+ zD2) or a control plasmid pEF-BOS-HA. Cells were pre-incubated for 4 hours with increasing concentrations of TTZ-1 (**A**) or TTZ-2 (**B**) or DMSO (as a control) before metabolic labelling with 100 μM C16:0-azide for another 4 hours. Cells were lysed and proteins having incorporated C16:0-azide were labelled by click chemistry with an alkyne-IR800 dye. Protein samples were then resolved by SDS PAGE and transferred to nitrocellulose membranes that were probed with an anti-GFP antibody (IR680). Representative images are shown for cells treated with TTZ-1 (**A**) or TTZ-2 (**B**). *Top panels*: click chemistry signal (Top panel, *Click (IR800)*) and GFP immunoblot (*Middle panel, GFP (IR680)*). Positions of the molecular mass markers (in kDa) are indicated on the right side of the membranes. *Bottom panels **A** and **B***. Data from 3 independent experiments performed in duplicate were quantified and the inhibition of GFP-SNAP25b S-acylation was determined as stated in *Experimental Procedures*. Graphs show mean +/- SEM of inhibition of normalised S-acylation (Click signal / GFP signal) in the presence of each compound. Data were analysed and fit to a nonlinear curve using the log(inhibitor) versus normalised response (variable slope) equation.

### Inhibition profile of TTZ-1 and -2 towards different zDHHC enzyme isoforms in cell-based S-acylation assays

We next investigated whether the identified compounds have any preferential activity towards zDHHC2 compared to other zDHHCs (Figure 6 and Table 1). In addition to zDHHC2, SNAP25 has been shown to be a substrate for S-acylation by zDHHC3, -7, -15 and -17 when co-expressed in HEK293T cells (6,45). We therefore treated cells co-expressing GFP-SNAP25 and each of these zDHHC enzymes with 25 μM of TTZ-1 and TTZ-2 and studied the subsequent effect on SNAP25 S-acylation. Figure 6A shows complete inhibition of zDHHC2 mediated SNAP25 S-acylation by TTZ-1 (113 +/- 5 %) and a very strong reduction (84 +/- 6 %) by TTZ-2. Importantly, the level of enzyme expression was unaffected by the compounds. Interestingly, the inhibitory effect of the compounds on zDHHC15-mediated S-acylation of SNAP25 was similar to that seen with zDHHC2 (Figure 6D). In contrast, S-acylation of SNAP25 by zDHHC3 and zDHHC7 was less inhibited: TTZ-1 had no significant effect on either enzyme in this assay, and TTZ-2 caused a 19 +/- 3 % reduction of SNAP25 S-acylation by zDHHC3 and had no significant effect on S-acylation by zDHHC7 (Figures 6B and 6C). Finally, both compounds showed an intermediate effect on zDHHC17 mediated S-acylation of SNAP25, with a ~ 35-40 % reduction (Figure 6E). Additionally, there was a slight but significant reduction in the level of zDHHC17 expressed in cells treated with TTZ-2. As any major differences in the expression levels of the different enzymes could in theory contribute to the apparent differential effects of TTZ-1 and TTZ-2 we directly compared the expression of zDHHC-2, -3, -7, -15 and -17 when transfected in HEK293T cells together with GFP-SNAP25. As shown in Figure 6F, none of the zDHHC enzymes was expressed to a higher level than HA-zDHHC2, suggesting the observed differential effects of TTZ-1 and -2 are not due to increased expression levels of a particular enzyme.

**Figure 6.**
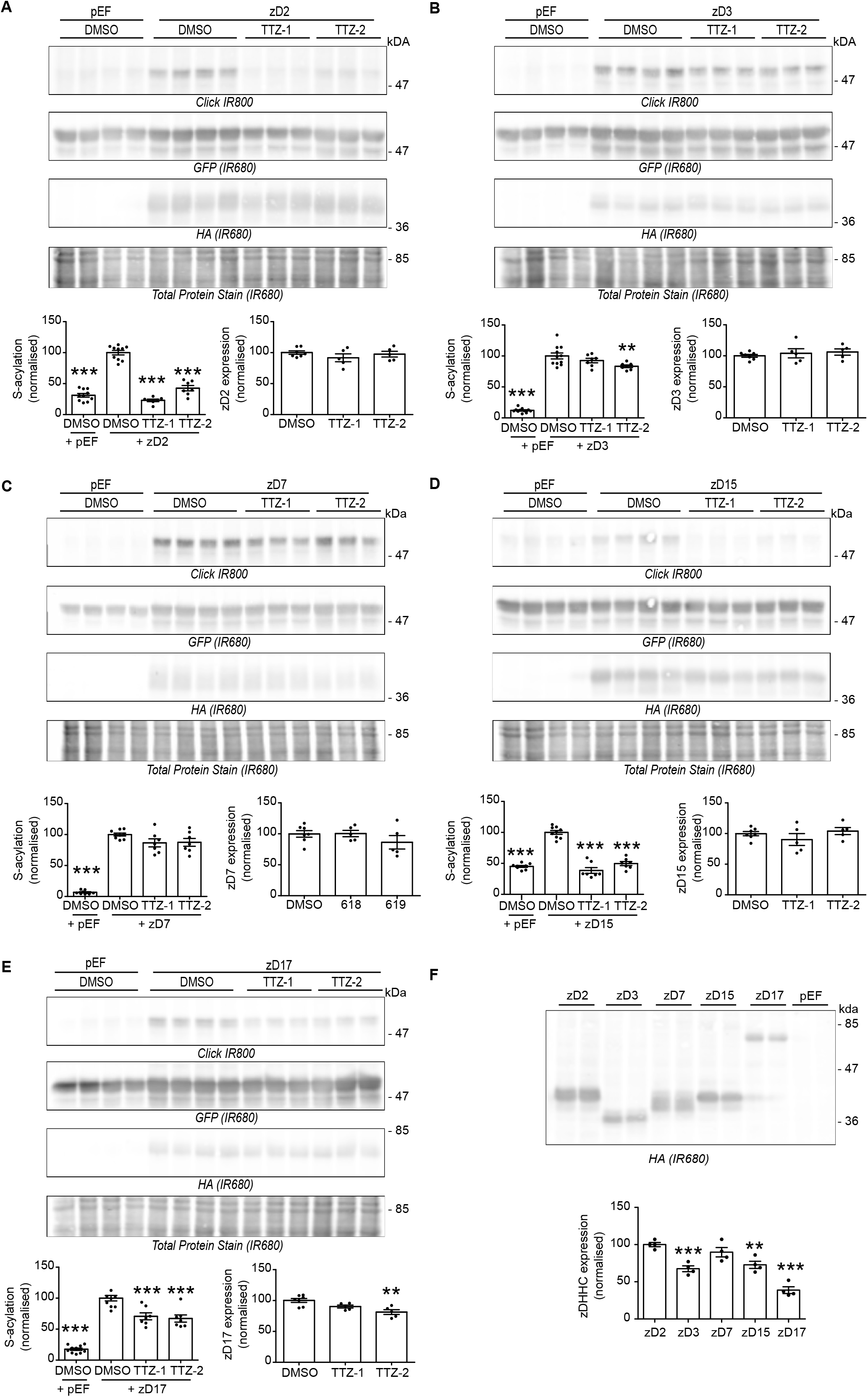
Activity of TTZ-1 and TTZ-2 against SNAP25 S-acylation by human zDHHC2, -3, -7, -15 and -17 expressed in mammalian cells. **A** to **E**. HEK293T cells were transfected with a plasmid encoding GFP-SNAP25b together with either a control plasmid pEF-BOS-HA (pEF) or a plasmid encoding either HA-tagged human zDHHC2 (zD2) (**A**), -3 (zD3) (**B**), -7 (zD7) (**C**), -15 (zD15) (**D**) or -17 (zD17) (**E**) for 24 hours. Cells were then preincubated for 4 hours with TTZ-1 or TTZ-2 (25 μM final) or DMSO control before metabolic labelling with 100 μM C16:0-azide for 4 hours. Cells were lysed and proteins having incorporated C16:0-azide were labelled by click chemistry with an alkyne-IR800 dye. Protein samples were then resolved by SDS PAGE and transferred to nitrocellulose membranes that were first stained with an infrared-680 total protein stain. Membranes were then destained and probed with an anti-HA antibody (IR680), followed by an anti-GFP antibody (IR680). Representative images are shown and panels are presented in the following order (from top to bottom): click chemistry signal (*Click (IR800)*), GFP and HA immunoblots (*IR680*) and Total Protein Stain (*IR680*). Positions of the molecular mass markers (in kDa) are indicated on the right side of the membranes. Three independent experiments were performed either in duplicate or triplicate and the quantification data were gathered on the graphs below each set of membranes. The graph on the left shows mean +/- SEM of normalised S-acylation of GFP-SNAP25b (Click signal / GFP signal) for 3 independent experiments. *Filled circles* represent individual samples. Statistical analysis (ANOVA) was performed to reveal whether there was a significant difference between the zDHHC enzyme mediated S-acylation of GFP SNAP25 treated with DMSO vs similar samples treated with TTZ-1 or -2 or vs cells that do not overexpress any zDHHC (+ pEF) (***, p<0.001; **, p<0.01). The graph on the right shows mean +/- SEM of normalised zDHHC expression (HA signal / Total Protein Stain signal) from 2 independent experiments. *Filled circles* represent individual samples. Statistical analysis (ANOVA) was performed to test whether there was a significant difference between the zDHHC expression of cells treated with DMSO as a control vs similar samples treated with TTZ-1 or TTZ-2 (**, p<0.01). **F.** Samples from **A**, **B**, **C**, **D** and **E** (expressing GFP-SNAP25 and HA-tagged zDHHC enzymes and treated with DMSO) were loaded on a SDS PAGE gel and transferred to a nitrocellulose membrane. The membrane was probed for the expression of the HA-tagged enzymes with an anti HA antibody (IR680). Positions of the molecular mass markers (in kDa) are indicated on the right side of the membranes. The graph represents the mean +/- SEM of normalised zDHHC expression (HA signal) from 2 independent experiments. *Filled circles* represent individual samples.

### Synthesis of TTZ-1 and TTZ-2 confirms the identity of the hit compounds and allows confirmation of biological results

To confirm the identity of the hit compounds and to allow further investigations to take place we prepared samples of TTZ-1 and -2 through the multi-step synthetic sequence outlined in Figure 7. One-pot sulfonation/bis-nitration of chlorobenzene **3** led to the sulfonate salt **4**. S_N_Ar reaction with aqueous ammonia followed by proto-desulfonation gave 2,6-dinotroaniline **6** in 36% overall yield for the initial sequence. Sandmeyer reaction of **6** under standard conditions gave 2,6-dinitrochlorobenzene **7** (57%) which was subjected to an efficient S_N_Ar reaction with hydroxide followed by partial reduction then reacted with 2-chloroacrylonitrile **10** to give the benzoxazine **11** in a poor, but acceptable yield of 15% after purification by silica gel chromatography. Reduction of the second nitro functionality led to the central scaffold of the target ligands **12** (70%) from which diversity could be introduced. Amide bond coupling followed by acid catalysed tetrazole formation gave TTZ-1 and TTZ-2. Compounds TTZ-1 and TTZ-2 were prepared in 12 synthetic steps and an overall yield of 1.9% and 1.5% respectively.

**Figure 7.**
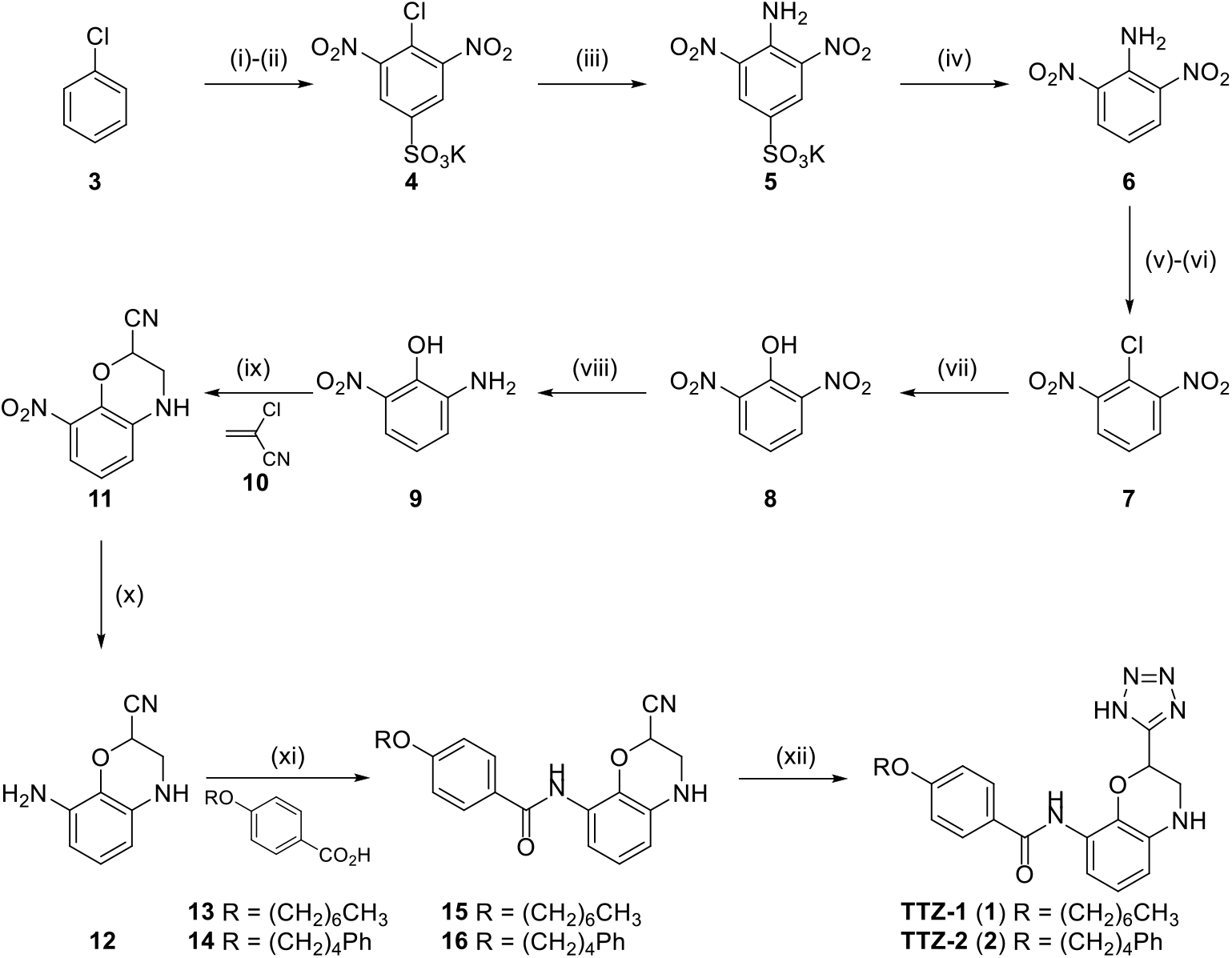
Synthesis of TTZ-1 (1) and TTZ-2 (2). *Reagents and conditions*: (i) H_2_SO_4(conc)_, 120 °C, 4 h. (ii) KNO_3_, 115 °C, 20 h. (iii) NH_3(aq)_, Δ, 1 h. (iv) H_2_SO_4(conc)_:H_2_O (1:1), Δ, 16 °C (36%, 4 steps). (v) NaNO_2_, H_2_SO_4_, 40 °C, 1 h. (vi) CuCl, HCl_(conc)_, 80 °C, 20 min (57%, 2 steps). (vii) KOH, H_2_O, Δ, 1 h (quant). (viii) 10% Pd/C, H_2_, MeOH, r.t., 1 h (88%). (xi) **10** (8.0 equiv.), Cs_2_CO_3_, PhMe, Δ, 4 h (15%). (x) 10% Pd/C, H_2_, MeOH, r.t., 1 h (70%). (xi) **13** (1 equiv.) or **14** (1 equiv.), HATU, DIPEA, DMF, r.t., 16 h (**15** 58%; **16** 45%). (xii) NaN_3_, NEt_3_·HCl, DMF, 140 °C, 2 h (**1** quant; **2** quant).

The newly synthesised compounds TTZ-1 and TTZ-2 were then tested against GFP-SNAP25 S-acylation in cell-based assays with results comparable to the original hit compounds (Supp Fig 2). We also tested the effects of these compounds on zDHHC2 S-acylation in HEK293T cells to confirm that the compounds affected autoacylation as seen in the original high-throughput screen using purified components (Figure 1). As shown in Figure 8, both compounds indeed significantly reduced the S-acylation of HA-zDHHC2 in HEK293T cells.

**Figure 8.**
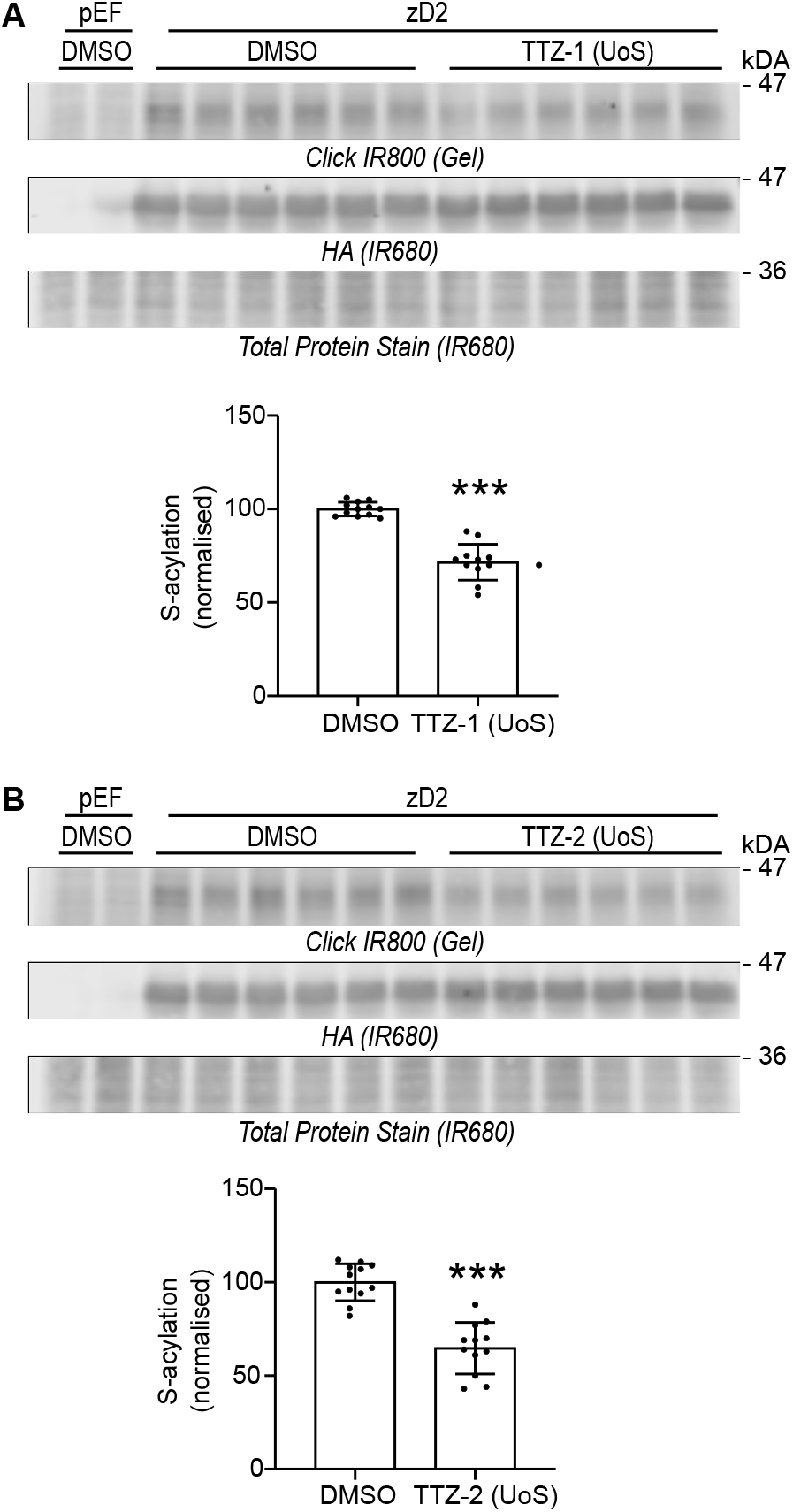
Activity of TTZ-1 (UoS) and TTZ-2 (UoS) against human zDHHC2 autoacylation expressed in mammalian cells. HEK293T cells were transfected with either the control plasmid pEF-BOS-HA (pEF) or a plasmid encoding HA-tagged human zDHHC2 (zD2) for 24 hours. Cells were then pre-incubated for 4 hours with UoS compounds TTZ-1 (UoS) (**A**) or TTZ-2 (UoS) (**B**) (25 μM final) or DMSO control before metabolic labelling with 100 μM C16:0-azide for 4 hours. Cells were lysed and proteins having incorporated C16:0-azide were labelled by click chemistry with an alkyne-IR800 dye. Protein samples were then resolved by SDS PAGE and the gel was scanned for the detection of the click chemistry signal in the infrared-800 channel. Gels were then transferred to nitrocellulose membranes that were first stained with an infrared-680 total protein stain. Membranes were then destained and probed with an anti-HA antibody (IR680). Representative images are shown and panels are presented in the following order (from top to bottom): click chemistry signal within the gel (*Click (IR800) (Gel)*), HA immunoblots (*IR680*) and Total Protein Stain (*IR680*). Positions of the molecular mass markers (in kDa) are indicated on the right side of the membranes. Two independent experiments were performed in sextuplicate and the quantification data were gathered on the graphs below each set of membranes. They show mean +/- SEM of normalised S-acylation of HA-zDHHC2 (Click signal / HA signal). *Filled circles* represent individual samples. Statistical analysis (Student’s t-test) reveals that there is a significant difference between the S-acylation of zDHHC2 in cells treated with DMSO vs similar samples treated with TTZ-1 or -2 (***, p<0.001).

## DISCUSSION

Despite the increasing recognition of the importance of S-acylation in physiological and pathological processes, progress in understanding the widespread functions of this post-translational modification is slowed by a dearth of available chemical modulators for use as tools compounds. Very few assays that would be amenable for HTS have been developed for the detection of S-acylation in the past 10 years. An early assay was based on the fluorescent detection of the released free CoA following S-acylation (46). The subsequent screen of a scaffold ranking library consisting of 68 unique scaffolds and 30 million unique structures successfully identified several Erf2 inhibitors (41). A click chemistry based HTS platform was also developed for the screening of Ras S-acylation inhibitors using membrane preparation as a source of enzyme, a biotinylated Ras peptide bound to streptavidin plates and alkyne palmitoyl-CoA as a lipid substrate. S-acylation was detected by the addition of azido-CalFluor488, a probe that is non-fluorescent until it participates in a click reaction with an alkyne entity (47). Although promising, no HTS result was subsequently reported with the use of this technique. The most recent report (Acyl-cLIP) detects the change in fluorescence polarisation of peptides (i.e. substrates) that follows the post translational attachment of lipid moieties and their subsequent binding to detergent micelles (48). A full HTS was thereafter performed to identify inhibitors of the activity of the N-palmitoyltransferase Hhat and will be reported in due course according to the authors. Acyl-cLIP was also tested for the detection of zDHHC3, 7 and 20 activities, showing that this assay can be adapted for the screening of zDHHC enzyme modulators (33,39,49).

Our focus here was to advance this area by: (i) developing new assay platforms for inhibitor discovery, and (ii) using these platforms to define novel chemical inhibitors of zDHHC enzymes. We showed that the developed high-throughput autoacylation assay was indeed a robust platform as it led to the identification of hits from a screen of >350,000 individual compounds that also inhibited enzyme autoacylation and substrate S-acylation in established cell-based assays. The compounds which emerged from this screen TTZ-1 and TTZ-2 both possess a 2,8-disubstituted 3,4-dihydro-2H-benzo[b][1,4]oxazine core. Common to both structures is a tetrazole head group, a well-established bioisostere for carboxylic acids. In addition, both compounds contain a hydrophobic chain at the 8-position of the heterocycle linked through an amide bond. The resemblance of the polar and non-polar components of these synthetic compounds to the natural fatty acid substrates for the zDHHC enzymes suggest that they may have a common binding site on the enzyme.

An important strength of the study is that the identified inhibitors displayed activity in multiple cell-free and cell-based assays, undertaken at different research sites. The compounds were indeed first identified for their inhibitory activity towards the auto-acylation of zDHHC2, as measured by TR-FRET. TR-FRET is a popular choice for HTS campaigns due to its relative simplicity and has been widely used for the screening of modulators of molecular interactions (protein-protein or protein-DNA) for example. TR-FRET has also been used to identify inhibitors of several enzymes such as kinases, heparanase, and ubiquitin ligases (50). This is to our knowledge the first report of the adaptation of TR-FRET for the detection of protein lipidation, and one of the largest screen of compounds modulating zDHHC mediated S-acylation. We have checked the specificity of the TR-FRET signal for S-acylation including using lipoparticles of inactive zDHHC2 mutant (C157S, Figure 1B). We also attempted to develop a TR-FRET assay for substrate acylation. However, with the materials we used in this publication (NBD-tagged palmitoyl-CoA, HA-zDHHC2, GST-SNAP25), we could not observe any TR-FRET signal on SNAP25 protein.

Following the identification of TTZ-1 and -2 in the HTS, the next step in the selection process was to screen their inhibitory activity towards substrate S-acylation in cell-free assays that used click chemistry for detection by either a newly-developed ELISA assay or by direct SDS-PAGE analysis. A click-ELISA assay was developed previously for the detection of Hhat activity (51) with the use of streptavidin coated plates for the attachment of a biotinylated substrate. S-acylation of the substrate was performed with purified enzyme and alkyne palmitoyl-CoA which was then linked to a FLAG tagged azide through copper catalysed click chemistry. A classic ELISA assay then followed for the detection of the FLAG epitope. In our hands the standard click chemistry reagent mixture impaired the binding of the GST tagged substrate to the glutathione plate and we therefore had to implement a less widely used click chemistry technique. Copper-free click chemistry involves the biorthogonal reaction between DBCO and an azide moiety and is carried out in the physiological buffer PBS, which did not disrupt the efficient binding of GST to the glutathione plate. Inconveniently, due to the reaction of DBCO with free cysteines the (non-acylated) residues had to be blocked with NEM prior to the click chemistry reaction. It should be noted that, in addition to the main distinction that these different assays measured either auto-acylation or substrate S-acylation, the TR-FRET and click chemistry assays also used different sources of zDHHC2 and different lipid substrates, highlighting the robustness of the generated data.

Finally, the compounds were tested for their ability to inhibit substrate S-acylation in cells expressing different zDHHC isoforms. It is interesting to note that the inhibition profile of TTZ-1 and -2 on SNAP25 S-acylation appeared to display some zDHHC isoform selectivity and were more potent against zDHHC2 and zDHHC15 *versus* zDHHC3 and zDHHC7, with zDHHC17 displaying intermediate levels of inhibition. We cautiously interpret these observations as indicating that the development of isoform-selective zDHHC inhibitors is a realistic possibility, perhaps due to subtle differences in the acyl-CoA binding pocket of zDHHC enzymes that affects lipid selectivity; this possibility will be explored in future work in this area. Supplementary figure 3 shows a phylogram of the relatedness of the human zDHHC family members based on the amino acid sequence of the 51-amino acid DHHC-CRD. As can be seen, zDHHC2 and zDHHC15 are closely related, which is consistent with the strong inhibition of both enzymes by TTZ-1 and -2. In contrast, less inhibition was seen towards zDHHC-3, -7 and -17, which are more distantly related to zDHHC-2/-15. We note that zDHHC3 and zDHHC7 are highly active isoforms (52) and therefore observed differences in inhibition profile might be related to differences, for example, in autoacylation turnover kinetics. To confirm isoform selectivity of these new S-acylation inhibitors will require detailed kinetic analyses using purified protein components and therefore we emphasise only the *potential* for these compounds to be further developed to enhance zDHHC enzyme isoform selectivity profiles. Nevertheless, the different effects of the compounds on different zDHHC enzymes does argue against their effects being indirect, for example, affecting fatty acid synthesis, uptake or conversion to acyl-CoA.

The further development of more potent and selective inhibitors of zDHHC15 could have utility for the treatment of induced autoimmune disease involving the constitutive activation of STING (Stimulator of Interferon Genes). STING can be S-acylated by zDHHC-3, -7 and -15, and the inhibition of STING S-acylation effectively reduces its activity, perhaps offering opportunities for the treatment of some inflammatory diseases (53). zDHHC2 inhibition could be of use for the management of several auto-immune disorders involving the lymphocyte-specific protein tyrosine kinase lck. S-acylation of this protein, which is essential for T cell activation, is mediated by zDHHC2 (54–56). There may also be potential for employing selective inhibitors of zDHHC enzymes for the treatment of some infectious diseases. Many viral proteins undergo S-acylation, and this is often key for efficient viral replication (57,58). Thus, acutely targeting the cellular enzymes that mediate S-acylation of essential viral proteins could interfere with viral infection. Indeed, the hemagglutinin of Influenza A was recently shown to be modified by zDHHC-2, -8, -15 and -20 (59). Similarly, the spike protein of SARS-CoV-2 can be S-acylated by zDHHC-2, -3, -6, -9, -11, -12, -20, -21 and -24 (42,60,61), and siRNA mediated knock down of zDHHC20 impairs virus infectivity (60). Another relevant example is the report that zDHHC2 is one of the major enzymes (together with zDHHC19) responsible for S-acylation of the RNA-binding protein nsP1 of the alphaviruses Chikungunya. Disrupting nsP1 S-acylation severely impaired viral replication (62). The development of highly selective zDHHC inhibitors may therefore offer scope for blocking S-acylation of viral proteins, whilst limiting effects on the S-acylation of host proteins. Alternatively, targeting S-acylation of host proteins such as CCR5 has been suggested as a potential strategy for the inhibition of HIV-1 infection (63). Recent data suggest that zDHHC-3, -7 and -15 are capable of S-acylating CCR5 when co-expressed in HEK293T cells, and that the inhibition of CCR5 S-acylation by two newly identified compounds is effective at reducing HIV-1 entry and replication in human macrophages. Interestingly, these compounds (cadmiun chloride and zinc pyrithione) are both zinc chelators and are proposed to act by binding directly to the zDHHC catalytic site of one or more of the enzymes identified; Zn^2+^ is known to bind to the DHHC-CRD of zDHHC enzymes and play an important structural role.

As shown in Supplementary figure 3, zDHHC20 also clusters phylogenetically with zDHHC2/15 (6,20,64) and it is therefore possible that the inhibitors identified here also target this enzyme. zDHHC20 has been implicated in the development of cancers resistant to EGFR inhibitor therapies (65–67). Interestingly, zDHHC20 mediates EGFR S-acylation and inhibits EGF signalling; silencing of zDHHC20 restores EGFR sensitivity and sensitizes cells to EGFR inhibitor-induced cell death, indicating that inhibition of zDHHC20 activity could function therapeutically in combination with EGFR inhibitors (66). Blocking zDHHC20-mediated EGFR S-acylation has also been shown to reduce PI3K signalling and MYC levels, and suppress cell growth *in vitro* and tumour growth in an *in vivo* model of KRAS-mutant lung adenocarcinoma (65).

Based on the results of the TR-FRET assays *in vitro* and of click chemistry *in cellulo*, the inhibitors described presumably target the first step of the enzymatic reaction, i.e. the auto-acylation of the enzyme, which makes them likely to inhibit the S-acylation of a broad range of substrates. Identifying inhibitors that are specific for a particular enzyme – substrate pair might be possible by targeting the interaction site between the two partners (30–32). With 23 human zDHHC enzymes and thousands of substrates, little is still known regarding the mode of interaction between enzyme and substrate (18,32,68). zDHHC-3, -5 and -7 possess a PDZ-binding C-terminal domain allowing for the specific recruitment of some of their substrates (18,68–71). In contrast, other substrates seem to display a PDZ-independent mode of binding to zDHHC5 (72,73), suggesting that compounds could be tailored to block specific enzyme-substrate pairs. For example zDHHC5 mediated palmitoylation of phospholemman (PLM), a small accessory subunit of the cardiac pump Na/K ATPase (NKA) is mediated through the recruitment of NKA by a juxtamembrane amphipathic α-helix of zDHHC5. Disrupting the interaction between NKA and zDHHC5 resulted in an inhibition of PLM palmitoylation and was achieved by the addition of a specifically designed cell penetrating peptide (74). There is also a growing understanding of how zDHHC17 recognizes and binds to its substrates. A key substrate recognition site between zDHHC17 and substrates is its N-terminal Ankyrin Repeat Domain (ARD), which recognizes a conserved sequence known as the zDABM in its substrates (18,75–79). Genistein has been recently identified as an inhibitor of MAP2K4 activity that disrupts the interaction between zDHHC17 ARD and MAP2K4, through direct binding to the ARD of zDHHC17; it would be worthwhile exploring the activity of Genistein against other zDHHC17 substrates (80). A substrate-targeted approach has also been successful recently with the identification of Ketoconazole as an inhibitor of the S-acylation and signalling of DLK (Dual Leucine-zipper Kinase) (81). Another potential angle of attack would be the interaction between zDHHCs and co-factors, such as the one between zDHHC9 and GCP16 (82), or zDHHC6 and selenoprotein K (83) for example.

The identification of small molecule inhibitors of zDHHC enzymes described herein provides valuable and much-needed tools to investigate S-acylation-dependent cellular pathways. These tools are significantly more drug-like in character and offer an important advance over non-selective inhibitors such as 2BP. It is envisaged that they will allow the acute effects of S-acylation disruption to be investigated without the need for longer-term zDHHC depletion (e.g. using RNAi or CRISPR technology). In addition, the apparent selectivity of these compounds towards zDHHC2 and zDHHC15 offers the potential to refine and develop their selectivity further through a detailed SAR study. Truly selective inhibitors of the zDHHC enzyme family would represent a major development for the field.

## EXPERIMENTAL PROCEDURES

### Cells

HEK293T cells (CRL-3216, ATCC) were grown at 37°C in a humidified atmosphere containing 5% CO_2_ in DMEM media (31966047, Gibco, Thermo Fisher Scientific) supplemented with 10% foetal bovine serum (11550356, Gibco, Thermo Fisher Scientific). Insect cells Sf9 (600100, Oxford Expression Technologies, UK) and Sf9 TriEx (71023-3, Novagen, Merck, UK) were grown at 28°C in a dry incubator without CO_2_, in ESF21 media (500300, Oxford Expression Technologies) and TriEx medium (71022, Novagen, Merck) respectively. All the components of the Expi293 MembranePro system (including the cells) were from Invitrogen (JPN).

### Antibodies

Goat GST antibody (27457701V, GE Healthcare, used at 1:1,000), NeutrAvidin-DyLight 800 (22853, Invitrogen, used at 1:5,000) and High Sensitivity NeutrAvidin-HRP (31030, Pierce, used at 1:20,000) were obtained from Thermo Fisher Scientific (UK). Rat HA antibody (Roche, clone 3F10, used at 1:1,000) was from Sigma (Poole, UK), mouse GFP antibody (Clontech, clone JL8, used at 1:4,000) was obtained from Takara (Saint-Germain-en-Laye, France). IR dye conjugated secondary antibodies were used at a dilution of 1:20,000 and purchased from LI-COR Biosciences (Cambridge, UK). Terbium cryptate conjugated anti HA antibody was from CisBio (Chiba, Japan).

### DNA plasmids

Human cDNA encoding zDHHC2 (NM_016353.4), -3 (NM_001135179), -7 (NM_017740.2), -15 (NM_144969.2) and -17 (NM_015336.2) were synthesised by GeneArt technologies (Thermo Fisher Scientific, UK) and sub-cloned in pEF-BOS-HA (6) in frame with the triple HA tag coding sequence at the N-terminal end.

The human cDNA encoding zDHHC2 was codon optimised for protein expression in insect cells and synthesized by GeneArt technology (Invitrogen). It was inserted with N and C-terminal tags (His6 and StrepII, respectively) in the baculovirus transfer vector pIEX/Bac3 (Merck Novagen #717263).

The validity of all constructs was verified by DNA sequencing (Dundee DNA Sequencing Service, UK).

### Fatty acids

C16:0-azide has been described previously (13). Palmitoyl-CoA-NBD (810705) was purchased from Avanti Polar Lipids. Azide-C16:0-CoA was synthesised using the method of Bishop and Haira (84).

### High throughput screen for the identification of zDHHC2 autoacylation inhibitors

The Expi293 MembranePro™ system (Invitrogen) was used to generate HA-tagged zDHHC2 lipoparticles according to manufacturer’s instructions. Briefly, 90 ml of Expi293 cells were co-transfected with 30 μg of pEF-BOS-HA-zDHHC2 plasmid and 90 μg of a plasmid encoding the lentiviral protein Gag. A transfection enhancer was added the next day. The supernatant of the cell culture (containing the zDHHC2 lipoparticles) was collected 72 hours post-transfection and mixed with the precipitation reagent provided in the kit for 1 or 2 days at 4 °C. The lipoparticles were then pelleted by centrifugation at 6,000 g for 30 minutes at 4 °C and resuspended in the buffer provided. The optimized protocol for the HTS was as follows: compounds were tested at a final concentration on 4.2 μM (2.5 nl) pre-mixed for a few minutes with 1.5 μl of HA-zDHHC2 lipoparticles (at 42 μg/ml) before the addition of 4.5 μl of a mixture of anti-HA Tb crypate (1:200) and NBD-palmitoyl-CoA (1 μM). Compounds were dispensed by Echo 555 (Beckman Coulter) and other reagents were dispensed by Multi Drop Combi (Thermo Fisher Scientific) The TR-FRET signal was read after a 15 minute incubation on a Biotek Synergy Neo apparatus (Agilent).

TR FRET ratio was calculated by Biotek Gen5 software (Agilent) using the equation: Acceptor signal/Donor signal*10000. The % inhibition was normalized by vehicle of WT enzyme (neutral control) and C157A inactive mutant (inhibitor control). Calculation of % inhibition and Quality control of whole campaign was made with Screener software (Genedata)

### Purification of recombinant proteins

The recombinant Baculovirus was obtained following manufacturer’s instructions (Oxford Technologies BaculoComplete kit #400100). Briefly, Sf9 cells (Oxford Expression Technologies #600100) were plated in 6-well plates and co-transfected with 1.2 μl of baculoFectin II, 100 ng of flashback Ultra DNA and 500 ng of transfer vector (pIEX-Bac3-hDHHC2 or the control vector expressing lacZ). Cells were incubated for 5 days at 28 °C before the supernatant (containing the recombinant baculovirus seed stock) was harvested and stored at 4 °C until required. Virus amplification was performed in Sf9 TriEX cells (Novagen / Merck #71023-3) grown in TriEx media (Novagen / Merck #71022). The media was removed from cells in T75 flasks (at ~ 50% confluence). 100 μl of the seed stock virus was diluted to 2 ml with medium and added to the cells. The virus was allowed to adsorb for 1 hour, with periodic rocking, before being removed. 15 ml of fresh medium was then added and the cells incubated for at least 5 days before the medium containing recombinant virus was harvested (this virus stock is the inoculum) and centrifuged at 1,000g for 20 minutes at 4 °C to remove broken cells. The recombinant viruses were titrated by serial dilution on 6-well plates of Sf9 cells and plaque titration.

Large scale production of zDHHC2 was performed in Sf9 TriEX cells. Medium was removed from 15 T75 flasks of TriEX cells and the recombinant virus added to the cells in a total volume of 2 ml and at a MOI of 2. The virus was allowed to adsorb at room temperature for 1 h, during which time the cells were occasionally gently rocked. The inoculum was then removed and replaced with 13 ml of fresh TriEX medium and incubated for 96 hours at 28 °C. The cells were then detached by tapping the flask, and collected by centrifugation at 1,000g for 5 minutes. 10 ml of PBS (pH 6.4) was added to the flask to collect the remaining cells, which were combined with the cell pellets and centrifuged again at 1,000g for 5 minutes. The cell pellets was then washed two further times in PBS (pH 6.4) and resuspended in a small volume of PBS supplemented with protease inhibitors (50X stock contains: Benzamidine HCl 800 μg/ml (Sigma #B65506), Aprotinin 500 μg/ml (Sigma #A1153), Leupeptin 500 μg/ml (Sigma #L2884), Pepstatin A 500 μg/ml (Sigma #P5318), PMSF 50 mM (Sigma #P7626)), pelleted and frozen at -80 °C until required.

Cell pellets were thawed and resuspended in lysis buffer (50mM Tris (Sigma #T6066) pH7.4, 200 mM NaCl (Sigma #31434), 10 %glycerol (Fisher #G/0600/17), 1% nDodecylβDMaltoside (DDM) (Thermo #89903), 1mM tris (2 carboxyethyl) phosphine hydrochloride (TCEP) (Sigma #C4706) and protease inhibitors (as above). 1 ml of lysis buffer was added per cell pellet (from 3 T75 flasks) and the cells disrupted further by 10 strokes of a Dounce homogeniser. The cell lysates were then rotated for 30 minutes at 4 °C on a wheel at a gentle speed and then clarified by ultracentrifugation at 100,000 g for 30 minutes at 4 °C. The lysates were then supplemented with 5-10 mM imidazole (Sigma #I5513), added to 1 ml of washed Ni^2+^-NTA agarose (Qiagen, #1018244) and rotated on a low speed wheel for at least 1h at 4 °C. The agarose was then pelleted by centrifugation at 2,000g for 5 minutes at 4°C and was washed 5 times with 10 ml of Wash Buffer (50 mM Tris pH7.4, 200 mM NaCl, 10% glycerol 0.2% DDM, 0.5 mM TCEP) containing 15 mM Imidazole. The washed agarose beads were resuspended in 500 μl of Elution Buffer 1 (50 mM Tris pH 7.4, 100 mM NaCl, 10% glycerol, 0.1% DDM, 0.25 mM TCEP, 200 mM Imidazole) and rotated on a wheel for 10 minutes at 4 °C. The agarose was then pelleted by centrifugation at 2,000g for 5 minutes at 4 °C. This elution step was repeated once. The agarose beads were then eluted 4 times with 500 μl of Elution Buffer 2 containing 500 mM Imidazole (50 mM Tris pH 7.4, 100 mM NaCl, 10% glycerol, 0.1% DDM, 0.25 mM TCEP, 500 mM Imidazole). The eluted samples were pooled (~ 3 ml) and dialysed twice (overnight and for 4 hours) against 500 ml of Elution Buffer without imidazole or TCEP. Purified proteins were analysed by SDS-PAGE and Instant Blue (10616474, Expedeon, ThermoFisher Scientific) staining. The concentration of the enzyme was determined by comparison with BSA standards and found to be between 1.5 and 3 μM depending on the batch.

The production and purification of GST and GST-SNAP25b has been described in (77). Briefly, *Escherichia coli* BL21(DE3)pLysS cells (#L1195, Promega) were transformed with either pGEX-KG (containing the GST cDNA) or pGEX-KG-rat SNAP25b (for the production of SNAP25b with a N-terminal GST tag) and selected with the appropriate antibiotics. Transformed cells were grown in a 1 litre culture of supermedia (150 mM NaCl, 1.5 % tryptone (Oxoid #LP0043), 2.5 % yeast extract (Oxoid #LP0021)) with shaking at 225 rpm (37 °C) and protein expression induced by the addition of 1 mM IPTG for 5-6 hours. Cells were pelleted and resuspended in PBS. The cells were lysed by subjecting to a freeze-thaw cycle at -80 °C, addition of 1 mg/ml lysozyme (Fluka #62971) and incubation for 30 minutes on ice. The cells were then sonicated, and cell debris and membranes removed by centrifugation at 20,000 xg for 60 minutes. The clarified lysates were incubated with glutathione sepharose (1 ml bed volume; GE Healthcare #17-0756-01), washed and eluted by the addition of 2 x 1.5 ml of 10 mM reduced L-glutathione (Sigma #T6066) in 50 mM Tris pH 8. Eluted proteins were dialysed overnight at 4°C against 5 l of PBS in a 3.5 MWCO slide-A-lyzer G2 cassette (#87723; Thermo). Purified proteins were analysed by SDS-PAGE and Instant Blue staining. Their concentration was estimated form the intensity of their corresponding bands as compared to the standard curve obtained with BSA standards that were run in parallel.

### In vitro substrate S-acylation followed by copper-free click chemistry

In an uncoated 96 well plate, 0.075 μM of purified zDHHC2 enzyme was incubated with 1 μM GST-tagged substrate protein and 1 μM Azide-C16:0-CoA at 25 °C for 45 minutes. The Assay Buffer consisted of 50 mM MES pH 6.4, 100 mM NaCl, 0.1% DDM, and 1 mM TCEP. Following the reaction, NEM was added to a final concentration of 50 mM and the plate incubated for 45-60 minutes at room temperature with gentle rocking. Following this, 10 μl of click reagent (Dibenzocyclooctyne biotin conjugate; Sigma #760749) diluted in PBS was added (5 μM final) and incubated at room temperature for one hour with rocking. 100 μl of PBS was added to each well of the uncoated plate to dilute the samples. For assessment of Ono compounds using this assay, the compounds were pre-incubated with enzyme for 15 minutes at room temperature prior to the addition of substrates. The concentration of compounds used refers to their final concentration in the S-acylation reaction and they were therefore added at a higher initial concentration (1.66x) for the 15 min pre-incubation period.

### Detection of in vitro substrate S-acylation by SDS PAGE and western blotting

Samples prepared as detailed above were supplemented with 4x SDS PAGE loading buffer (containing 100 mM DTT). They were heated at 95°C for 5 min and loaded on a SDS PAGE gel before being transferred to a nitrocellulose membrane. The membrane was incubated with NeutrAvidin-800 and a goat anti-GST antibody and then an anti-goat-680 antibody before being imaged on a LI-COR Odyssey infrared scanner.

### Measurement of in vitro substrate S-acylation by ELISA

Glutathione-coated plates were activated (according to manufacturer instructions; Pierce Glutathione Coated Black 96 well plates Thermo #15340) with 2 brief washes with 200 μl of PBS 0.05% v/v Tween-20. 80 μl of PBS was then added to each well and 20 μl of each sample (prepared as above) added (in duplicate) and incubated at room temperature for 1-2h on a rocking platform. The plate was then washed several times with 200 μl of PBS-0.05% Tween-20. 100 μl of NeutrAvidin-HRP (Pierce High Sensitivity NeutrAvidin HRP, ThermoFisher Scientific, #31030) diluted at 1:20,000 in PBS 0.025% Tween-20 was added to each well and incubated at room temperature for 45 minutes on a rocking platform. The plate was then washed several times with 250 μl of PBS-0.05% Tween-20 and incubated with 100 μl of SuperSignal ELISA Pico chemiluminescent substrate (Thermo Scientific #37069). The resulting signal was read 10 minutes post addition of the substrate For quantification, the average value of GST-SNAP25 samples (without zDHHC2) was subtracted from all raw values. These values were then normalised to the value of the DMSO control sample containing GST-SNAP25 and zDHHC2, providing a percentage of activity remaining after treatment with inhibitors. The percentage inhibition was then calculated as (100 – percentage activity remaining). Data were analysed and fitted to a non-linear curve using the log(inhibitor) *versus* normalised response (variable slope) equation (GraphPad software).

### Assessment of the activity of compounds on S-acylation in mammalian cells

HEK293T cells were plated on poly-D-lysine coated 24-well plates (356413, Corning BioCoat, VWR, UK) and transfected with 2.4 μg of pEF-zDHHC2 plasmid (or the control empty plasmid pEF-BOS-HA) for the measurement of zDHHC2 auto-acylation and 4.8 μl of Lipofectamine 2000 (Invitrogen, UK). For substrate acylation cells were transfected with 1.6 μg of pEF-zDHHC plasmid (or the control empty plasmid pEF-BOS-HA) together with 0.8 μg of EGFP-SNAP25b plasmid and 4.8 μl of Lipofectamine 2000. Twenty-four h post-transfection, cells were washed once with PBS and preincubated with ONO or UoS compounds diluted in 250 μl of serum-free DMEM for 4 h. S-acylation labelling was then started by the addition of 50 μl of 600 μM C16:0-azide (final concentration 100 μM) diluted in serum-free DMEM supplemented with 1 mg/ml fatty-acid free BSA (Sigma, A7030). Cells were labelled for 4 h, washed once with PBS then lysed on ice in 100 μl of 50 mM Tris pH8 containing 0.5% SDS and protease inhibitors (Sigma, P8340). Conjugation of IR800 alkyne dye to C16:0-azide was carried out for 1 h at room temperature with end-over-end rotation by adding an equal volume of freshly prepared click chemistry reaction mixture containing the following: 5 μM IRDye 800CW alkyne (929-60002, Li-COR, UK), 4 mM CuSO_4_ (451657, Sigma, UK), 400 μM Tris[(1-benzyl-1*H*-1,2,3-triazol-4-yl)methyl]amine (678937, Sigma, UK) and 8 mM ascorbic acid (A15613, Alpha Aesar, UK) in dH_2_O. 67 μl of 4x SDS-PAGE sample buffer containing 100 mM DTT was then added to the 200 μl sample. Samples were heated at 95°C for 5 min and 20 μl was resolved by SDS-PAGE and transferred to nitrocelullose membrane for immunoblot analysis. Membranes were first stained with Revert 700 total protein stain kit (926-11016, LI-COR), imaged, destained and then analysed by immunoblotting. Signals were quantified with ImageStudio software (LI-COR). S-acylation was calculated as the ratio (in %) between click signal (IR800) and GFP signal (IR680) and normalised to control conditions on the same membrane (cells expressing GFP-SNAP25 + zDHHC and treated with DMSO). The expression of the various zDHHCs was also quantified as a ratio between the HA signal (IR680) and the Total Protein Stain (IR680) and normalised to control conditions on the same membrane (cells expressing GFP-SNAP25 = zDHHC and treated with DMSO). For the determination of the *in cell* IC_50_ the % of S-acylation inhibition was calculated as (100 - % of S-acylation). Data were analysed and fitted to a non-linear curve using the log(inhibitor) versus normalised response (variable slope) equation (GraphPad software).

### Phylogenic analysis of ZDHHC

The following human ZDHHC protein sequences were retrieved from the NCBI database: ZDHHC1 (NP_001310556.1), ZDHHC2 (NP_057437.1), ZDHHC3 (NP_001128651.1), ZDHHC4 (NP_001358225.1), ZDHHC5 (NP_056272.2), ZDDHC6 (NP_071939.1), ZDHHC7 (NP_060210.2), ZDHHC8 (NP_037505.1), ZDHHC9 (NP_057116.2), ZDHHC11 (NP_079062.1), ZDHHC12 (NP_001304944.2), ZDHHC13 (NP_061901.2), ZDHHC14 (NP_078906.2), ZDHHC15 (NP_659406.1), ZDHHC16 (NP_115703.2), ZDHHC17 (NP_056151.2), ZDHHC18 (NP_115659.1), ZDHHC19 (NP_001034706.1), ZDHHC20 (NP_694983.2), ZDHHC21 (NP_001341047.1), ZDHHC22 (NP_777636.2), ZDHHC23 (NP_001307395.1) and ZDHHC24 (NP_997223.1). They were analysed on www.phylogeny.fr. Sequences corresponding to the 51 amino-acid DHHC Cysteine-Rich Domain were aligned with MUSCLE, phylogenic analysis was performed with PhyML and the tree was rendered with TreeDyn.

### Synthesis of TTZ-1 and TTZ-2

#### 2.6-Dinitroaniline (6)

(85) Chlorobenzene **1** (20.0 mL, 0.20 mol) was added to concentrated sulfuric acid (150 mL) and the solution was heated at 120 °C for 4 h. After cooling the reaction to room temperature, KNO_3_ (68.0 g, 0.6 mol) was added in 4 portions while stirring. The reaction was then heated at 115 °C for 20 h while stirring. While still hot, the solution was poured onto 1 kg of ice, once all the ice was melted the suspension was filtered and pressed dry. The solid was then dissolved in hot distilled water (240 mL), most of insoluble solids were removed by decantation and filtration of the solution while still hot. The product was then left to crystallise, and filtered to give a pale-yellow solid which was dissolved in aqueous ammonia (180 mL) and water (180 mL). The mixture was left stirring at reflux for 1 h. The solution was cooled down to 4 °C for 12 h and the solid was filtered under suction to give an orange solid that was dissolved in 100 mL of concentrated sulfuric acid and 100 mL of water. The solution was heated at reflux for 16 h. While hot, the solution was poured onto 1 kg of cracked ice. When the ice had melted, the solid was filtered under vacuum to give the *title compound* **6** as a yellow solid (12.9 g, 36%). mp: 138 °C (Lit: 137–142 °C) (85); LRMS (ES + APCI): *m/z* calc. 183.1, found 182.1 [M-H]^-^; *υ*_max_ (thin film, cm^-1^) 3462, 3345, 3107, 1636, 1513; ^1^H NMR (400 MHz, DMSO-*d*_6_) δ 8.49 (d, *J* = 8.3 Hz, 2H), 8.39 (s, 2H), 6.84 (t, *J* = 8.3 Hz, 1H); ^13^C NMR (101 MHz, DMSO-*d*_6_) δ 140.8, 134.9, 134.0, 113.8.

#### 2-Chloro-1,3-dinitrobenzene (7)

(86). Sodium nitrite (1.09 g, 15.8 mmol) was added to concentrated sulfuric acid (11 mL), and the resulting solution was heated at 40 °C for 20 min to ensure that all the solid was dissolved, then a solution of 2,6-dinitroaniline **6** (2.63 g, 14.4 mmol) in hot acetic acid (27 mL) was added at such a rate that the temperature was kept under 40 °C. After the addition was complete, the reaction was heated at 40 °C for 1 h. Copper(I) chloride (2.1 g, 21.2 mmol) was added to ice cold hydrochloric acid (27 mL), then the diazonium salt solution was added so that the effervescence was kept under control. After the addition was finished the reaction was heated at 80 °C until the effervescence stopped. The reaction mixture was cooled to 4 °C for 12 h and then the solid was filtered under vacuum to give the *title compound* **7** as a yellow solid (1.65 g, 57%). mp: 87 °C (Lit: 86–87 °C) (86); *υ*_max_ (thin film, cm^-1^) 3103, 1701, 1528; ^1^H NMR (400 MHz, Acetone-*d*6) δ 8.33 (d, *J* = 8.2 Hz, 2H), 7.93 (t, *J* = 8.2 Hz, 1H), 6.01 (bs, 1H); ^13^C NMR (101 MHz, Acetone-*d*6) δ 150.5, 130.6, 129.3, 119.7.

#### 2.6-Dinitrophenol (8)

(87). 2-Chloro-1,3-dinitrobenzene **7** (1.65 g, 8.14 mmol) and potassium hydroxide (1.37 g, 16.3 mmol) were added to water (30 mL) and the solution was heated to reflux for 1 h. The hot solution was added to ice cold water (30 mL) and the pH was adjusted to 5 through the addition of concentrated HCl. The solution was extracted with EtOAc (25 mL × 3), the combined organic phase was dried over anhydrous magnesium sulfate and the solvent was removed under reduced pressure to give the *title compound* **8** as a red solid (1.50 g, quant). mp: 63 °C (Lit: 63.5 °C) (87); LRMS (ES + APCI): *m/z* calc. 184.1, found 183.1 [M-H]^-^; *υ*_max_ (thin film, cm^-1^) 3215, 3096, 1767, 1932, 1809, 1692, 1623, 1534; ^1^H NMR (400 MHz, Acetone-*d*6) δ 8.35 (d, *J* = 8.3 Hz, 2H), 7.20 (t, *J* = 8.3 Hz, 1H); ^13^C NMR (101 MHz, Acetone-*d*_6_) δ 150.3, 139.8, 131.8, 118.8.

#### 2-Amino-6-nitrophenol (9)

(88). Hydrogen was bubbled through a nitrogen purged solution of 2,6-dinitrophenol **8** (1.5 g, 8.15 mmol) and 10% palladium on activated carbon (150 mg, 1.60 mmol) in methanol (20 mL) for 15 min. The reaction mixture was then left stirring under a hydrogen atmosphere for 45 min. The reaction was monitored by TLC. After the starting material was consumed the reaction was filtered through Celite and eluted with acetone (50 mL) and the solvent was removed *in vacuo*. The crude was purified by flash chromatography (toluene:dichloromethane 9:1) to give the *title compound* **9** as a red solid (1.1 g, 88%). mp: 110 °C; LRMS (ES + APCI): *m/z* calc. 154.1, found 155.1 [M+H]^+^; *υ*_max_ (thin film, cm^-1^) 3481, 3385, 3223, 3103, 2921, 2852, 1538; ^1^H NMR (500 MHz, Chloroform-d) δ 10.73 (s, 1H), 7.48 (dd, *J* = 8.6, 1.5 Hz, 1H), 6.95 (dd, *J* = 7.7, 1.5 Hz, 1H), 6.79 (dd, *J* = 8.6, 7.7 Hz, 1H), 4.09 (s, 2H); ^13^C NMR (101 MHz, Chloroform-d) δ 143.4, 137.9, 133.8, 120.5, 119.9, 113.6.

#### 8-Nitro-3,4-dihydro-2*H*-benzo[*b*][1,4]oxazine-2-carbonitrile (11)

2-Amino-6-nitrophenol **9** (500 mg, 3.24 mmol) and caesium carbonate (2.11 g, 6.48 mmol) were dissolved in toluene (18 mL) and heated at 50 °C for 30 min. Then, 2-chloroacrylonitrile **10** (2.0 mL, 25.95 mmol) was added and the reaction was heated at 110 °C in a sealed vial for 4 h. After the reaction was completed, the solvent was removed under reduced pressure to give a brown solid which was filtered through a Celite plug and eluted with acetone. The solvent was removed *in vacuo* to give a brown solid. The crude product was purified on silica gel (toluene:EtOAc 8:2) to give the *title compound* **11** as a brown solid (100 mg, 15%). LRMS (ES + APCI): *m/z* calc. 205.2, found 204.0 [M-H]^-^; *υ*_max_ (thin film, cm^-1^) 3392, 3111, 2927, 1780, 1688, 1616, 1532; ^1^H NMR (500 MHz, Acetone-d6) δ 7.17 (dd, *J* = 8.1, 1.7 Hz, 1H), 7.05 (dd, *J* = 8.1, 1.7 Hz, 1H), 6.99 (at, *J* = 8.1 Hz, 1H), 6.15 (bs, 1H), 5.63–5.60 (m, 1H), 3.89–3.75 (m, 2H); ^13^C NMR (101 MHz, Acetone) δ 141.2, 136.1, 134.3, 123.1, 120.0, 116.8, 114.1, 63.2, 43.0.

#### 8-Amino-3,4-dihydro-2*H*-benzo[*b*][1,4]oxazine-2-carbonitrile (12)

8-Nitro-3,4-dihydro-2*H*-benzo[*δ*][1,4]oxazine-2-carbonitrile **11** (50 mg, 0.24 mmol) and 10% palladium on activated carbon (5 mg, 10 mol%) were dissolved in methanol (5 mL), and the solution was purged with hydrogen for 15 min and left stirring under a hydrogen atmosphere for 45 min. The solution was filtered through Celite, eluted with acetone and the solvent was removed under vacuum to give the *title compound* **12** as a brown oil (29 mg, 70%). LRMS (ES + APCI): *m/z* calc. 175.2, found 176.1 [M+H]^+^; *υ*_max_ (thin film, cm^-1^) 3370, 2966, 2863, 2251, 1623, 1498; ^1^H NMR (400 MHz, Chloroform-d) δ 6.68 (at, *J* = 8.0 Hz, 1H), 6.22 (dd, *J* = 8.0, 1.4 Hz, 1H), 6.12 (dd, *J* = 8.0, 1.4 Hz, 1H), 5.15–5.10 (m, 1H), 3.85 (bs, 1H), 3.74 (bs, 2H), 3.65–3.60 (m, 2H);^13^C NMR (101 MHz, CDCl_3_) δ 136.3, 132.0, 129.2, 123.1, 116.4, 107.2, 106.5, 62.7, 43.9.

#### *N*-(2-Cyano-3,4-dihydro-2*H*-benzo[*b*][1,4]oxazin-8-yl)-4-(heptyloxy)benzamide (15)

4-(Heptyloxy)benzoic acid **13** (5 mg, 0.02 mmol), HATU (8 mg, 0.02 mmol) were dissolved in DMF (1 mL) and *N,N*-diisopropylethylamine (11 μL, 0.06 mmol) was added, and the resulting solution was left stirring for 45 min. A solution of compound **12** (4 mg, 0.02 mmol) in DMF (100 μL) was added to the acid solution, and it was left stirring at room temperature for 16 h. An aqueous solution of 5% lithium chloride (30 mL) was added and the aqueous phase was extracted with EtOAc (20 mL × 3), the organic fractions were combined and dried over anhydrous MgSO_4_, the organic solution was filtered and the solvent was removed by rotatory evaporation. The crude was purified by flash chromatography (PhMe:EtOAc 8:2) to give the *title compound* **15** as a brown oil (6 mg, 58%). LRMS (ES + APCI): *m/z* calc. 393.2, found 394.2 [M+H]^+^; *υ*_max_ (thin film, cm^-1^) 3381, 2956, 2932, 2859, 2552, 1662, 1610, 1506;^1^H NMR (400 MHz, Acetone-d6) δ 8.68 (bs, 1H), 7.96–7.92 (m, 2H), 7.71 (dd, J = 8.1, 1.5 Hz, 1H), 7.09–7.01 (m, 2H), 6.82 (at, J = 8.1 Hz, 1H), 6.53 (dd, J = 8.1, 1.5 Hz, 1H), 5.58– 5.51 (m, 2H), 4.09 (t, J = 6.5 Hz, 2H), 3.79–3.65 (m, 2H), 1.86–1.76 (m, 2H), 1.55–1.26 (m, 8H), 0.96– 0.83 (m, 3H); ^13^C NMR (101 MHz, Acetone) δ 165.0, 163.0, 134.0, 131.6, 130.0, 129.2, 128.1, 123.0, 117.5, 115.1, 111.8, 68.9, 63.5, 43.9, 32.6, 26.7, 23.3, 14.3 3 carbons missing.

#### *N*-(2-Cyano-3,4-dihydro-2*H*-benzo[*b*][1,4]oxazin-8-yl)-4-(4-phenylbutoxy)benzamide (16)

4-(Benzyloxy)benzoic acid **14** (6 mg, 0.02 mmol), HATU (8 mg, 0.21 mmol) were dissolved in DMF (1 mL) and *N,N*-diisopropylethylamine (11 μL, 0.06 mmol) was added, and the resulting solution was left stirring for 45 min. A solution of compound **12** (4 mg, 0.02 mmol) in DMF (100 μL) was added to the acid solution, and it was left stirring at room temperature for 16 h. A solution of 5% lithium chloride (30 mL) was added and the aqueous phase was extracted with EtOAc (15 mL × 3), the organic fractions were combined and dried over anhydrous MgSO_4_, the organic solution was filtered and the solvent was removed by rotatory evaporation. The crude was purified by flash chromatography (PhMe:EtOAc 8:2) to give the *title compound* **16** as a brown oil (4 mg, 45%). LRMS (ES + APCI): *m/z* calc. 427.2, found [M+H]^+^; *υ*_max_ (thin film, cm^-1^) 3375, 3087, 3062, 3029, 2940, 2863, 2543, 1660, 1608; ^1^H NMR (400 MHz, Acetone-d6) δ 8.68 (bs, 1H), 7.97–7.91 (m, 2H), 7.72 (dd, J = 8.1, 1.5 Hz, 1H), 7.32–7.22 (m, 4H), 7.20–7.14 (m, 1H), 7.06–7.01 (m, 2H), 6.82 (at, J = 8.1 Hz, 1H), 6.53 (dd, J = 8.1, 1.5 Hz, 1H), 5.55–5.51 (m, 2H), 4.15–4.09 (m, 2H), 3.77–3.64 (m, 2H), 2.71 (t, J = 7.0 Hz, 2H), 1.91–1.72 (m, 4H); ^13^C NMR(101 MHz, Acetone) δ 165.0, 162.9, 143.2, 134.0, 131.6, 130.0, 129.2, 129.1, 128.1, 126.6, 123.0, 117.5, 115.1, 111.9, 68.7, 63.5, 43.9, 36.1, 28.7 3 carbons missing.

#### *N*-(2-(1H-Tetrazol-5-yl)-3,4-dihydro-2H-benzo[b][1,4]oxazin-8-yl)-4-(heptyloxy)benzamide

(TTZ-1) (**1**). Compound **15** (6 mg, 0.018 mmol) was dissolved in DMF (100 μL) then sodium azide (4 mg, 0.056 mmol) and triethylammonium chloride (8 mg, 0.056 mmol) were added and the vial was sealed and heated at 140 °C for 2 h. The contents of the vial were left to cool down to room temperature and a solution of 5% lithium chloride in water (30 mL) was added, the aqueous phase was extracted with ether (15 mL × 3), the organic fractions were combined and the solvent was removed *in vacuo* to give the *title compound* TTZ-1 (**1**) as a brown oil (7 mg, quant). LRMS (ES + APCI): *m/z* calc. 436.2, found 435.1 [M-H]^-^; *υ*_max_ (thin film, cm^-1^) 3325, 3271, 2951, 2928, 2874, 2859, 2735, 2617, 2470, 1640; ^1^H NMR (400 MHz, Acetone-d6) δ 9.49 (bs, 1H), 8.11–8.04 (m, 2H), 7.12–7.03 (m, 2H), 6.84 (dd, J = 8.0, 1.5 Hz, 1H), 6.75 (at, J = 8.0 Hz, 1H), 6.52 (dd, J = 8.0, 1.5 Hz, 1H), 5.89 (at, J = 3.0 Hz, 1H), 5.43 (bs, 1H), 4.11 (t, J = 6.5 Hz, 2H), 4.03–3.78 (m, 2H), 1.90–1.74 (m, 2H), 1.53–1.45 (m, 2H).1.59–1.20 (m, 6H), 0.95–0.82 (m, 3H); ^13^C NMR (101 MHz, Acetone-d6) δ 167.5, 163.4, 156.7, 136.1, 135.2, 130.7, 127.1, 126.7, 122.6, 115.7, 115.2, 114.3, 69.2, 69.0, 43.5, 32.5, 29.9, 26.7, 23.3, 14.3 1 carbon missing.

#### *N*-(2-(1*H*-Tetrazol-5-yl)-3,4-dihydro-2*H*-benzo[*b*][1,4]oxazin-8-yl)-4-(4-phenylbutoxy)benzamide

(TTZ-2) (**2**). Compound **16** (4 mg, 0.011 mmol) was dissolved in DMF (100 μL) then sodium azide (3 mg, 0.046 mmol) and triethylammonium chloride (3 mg, 0.022 mmol) were added and the vial was sealed and heated at 140 °C for 2 h. The contents of the vial were left to cool down to room temperature and a solution of 5% lithium chloride in water (30 mL) was added, the aqueous phase was extracted with ether (15 mL × 3), the organic fractions were combined and the solvent was removed *in vacuo* to give the *title compound* TTZ-2 (**2**) as a brown oil (5 mg, quant). LRMS (ES + APCI): *m/z* calc. 470.2, found 469.1 [M-H]^-^; *υ*_max_ (thin film, cm^-1^) 3331, 3027, 2928, 2858, 1701, 1638, 1608; ^1^H NMR (400 MHz, Acetone-d6) δ 9.49 b(s, 1H), 8.11–8.04 (m, 2H), 7.32–7.14 (m, 5H), 7.11– 7.04 (m, 2H), 6.84 (dd, J = 8.0, 1.6 Hz, 1H), 6.75 (at, J = 8.0 Hz, 1H), 6.52 (dd, J = 8.0, 1.6 Hz, 1H), 5.89 (at, J = 3.0 Hz, 1H), 5.43 (bs, 1H), 4.21–4.07 (m, 2H), 4.03–3.74 (m, 2H), 2.75–2.68 (m, 2H), 1.89–1.77 (m, 4H); ^13^C NMR (101 MHz, Acetone-d6) δ 168.1, 168.0, 167.5, 163.4, 151.8, 143.2, 130.7, 129.3, 129.2, 126.6, 122.5, 115.7, 115.2, 114.3, 69.2, 68.8, 55.5, 43.5, 36.1, 28.7 2 carbons missing.

## CRediT author statement

**Christine Salaun**: Conceptualization, Methodology, Investigation, Formal analysis, Writing-Original draft, Visualisation; **Hiroya Takizawa:** Conceptualization, Methodology, Investigation, Formal analysis, Writing-Review and editing, Visualisation; **Alex Galindo**: Methodology, Resources, Investigation, Writing-Original draft; **Kevin Munro**: Methodology, Resources, Writing-Review and editing; **Jayde McLellan**: Methodology, Resources, Writing-Review and editing; **Isamu Sugimoto**: Methodology, Resources, Writing-Review and editing; **Tomotaka Okino**: Conceptualization, Writing-Review and editing, Supervision; **Nicholas Tomkinson**: Conceptualization, Writing-Review and editing, Supervision, **Luke Chamberlain**: Conceptualization, Writing-Review and editing, Supervision.

## ACKNOWLEDGEMENTS

This work was funded by Ono Pharmaceutical Co. Ltd. We are grateful to Dr Maria Sanchez-Perez for contribution to establishing this collaborative project.

## DATA AVAILABILITY

All data is contained within the paper. Copies of H NMR and C NMR spectra are available as Supplementary data.

## SUPPLEMENTARY FIGURE LEGENDS

**Supplementary figure 1. Known Inhibitors of DHHC enzymes.**

**Supplementary figure 2. Activity of UoS-TTZ-1, UoS-TTZ-2 and TTZ-2 against SNAP25 S-acylation by human zDHHC2, -3, -7, -15 and -17 expressed in mammalian cells**

HEK293T cells were transfected with a plasmid encoding GFP-SNAP25b together with either a control plasmid pEF-BOS-HA (pEF) or a plasmid encoding either HA-tagged human zDHHC2 (zD2), -3 (zD3), -7 (zD7), -15 (zD15) or -17 (zD17) for 24 hours. Cells were then pre-incubated for 4 hours with compounds TTZ-1 and -2 (UoS), TTZ-2 (ONO) (25 μM final) or DMSO control before metabolic labelling with 100 μM C16:0-azide for 4 hours. Cells were lysed and proteins having incorporated C16:0-azide were labelled by click chemistry with an alkyne-IR800 dye. Protein samples were then resolved by SDS PAGE gel and transferred to nitrocellulose membranes that were first stained with an infrared-680 total protein stain. Membranes were then destained and probed with an anti-HA antibody (IR680), followed by an anti GFP antibody (IR680). Two independent experiments were performed either in duplicate (for TTZ-2 ONO) or triplicate (for TTZ-1 and -2 UoS) and the quantification data were gathered on the graphs showing mean +/- SEM of normalised S-acylation of GFP-SNAP25b (Click signal / GFP signal). *Filled circles* represent individual samples. Statistical analysis (ANOVA) was performed to reveal whether there was a significant difference between the zDHHC enzyme mediated S-acylation of GFP SNAP25 treated with DMSO vs similar samples treated with the compounds or vs cells that do not overexpress any zDHHC (+ pEF) (***, p<0.001; *, p<0.05).

**Supplementary Figure 3. Phylogenic tree of human zDHHCs based on 51 amino-acid DHHC domain.** Sequences were aligned with MUSCLE and phylogenic analysis was performed with PhyML.

