## supplementary data for "Development of a novel high-throughput screen for the identification of new inhibitors of protein S-acylation"

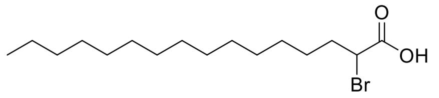

**2-bromopalmitic acid (2-BP)**

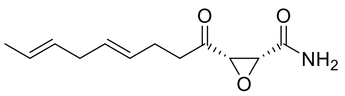

**cerulenin**

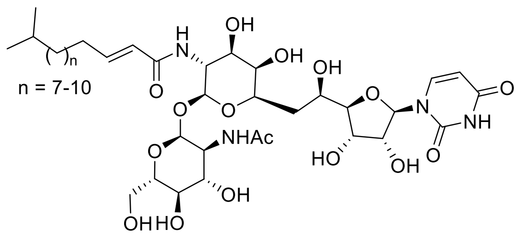

**tunicamycin**

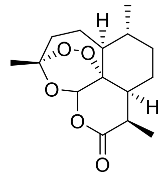

**artemisinin**

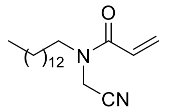

***N*-(cyanomethyl)-*N*-myracrylamide (CMA)**

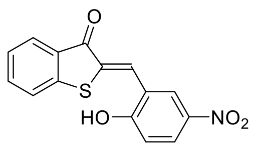

**Compound V**

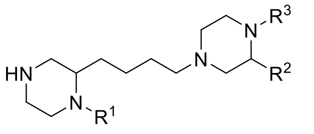

**bis-cyclic piperazine backbone**

### Supplementary Figure 1

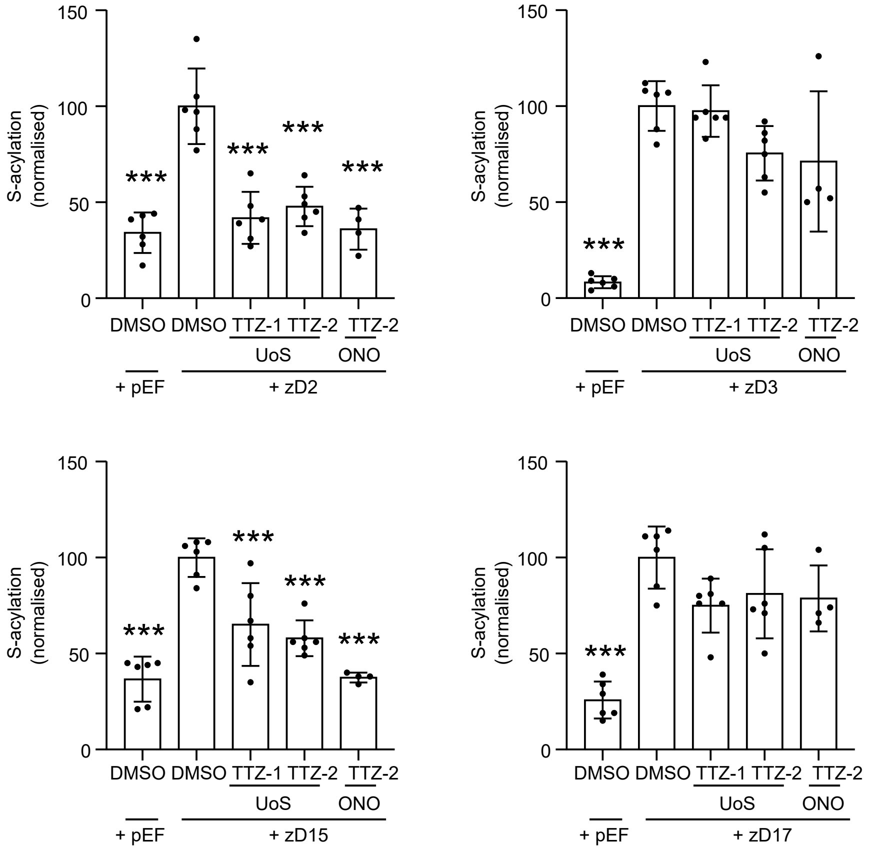


Supplementary Figure 2

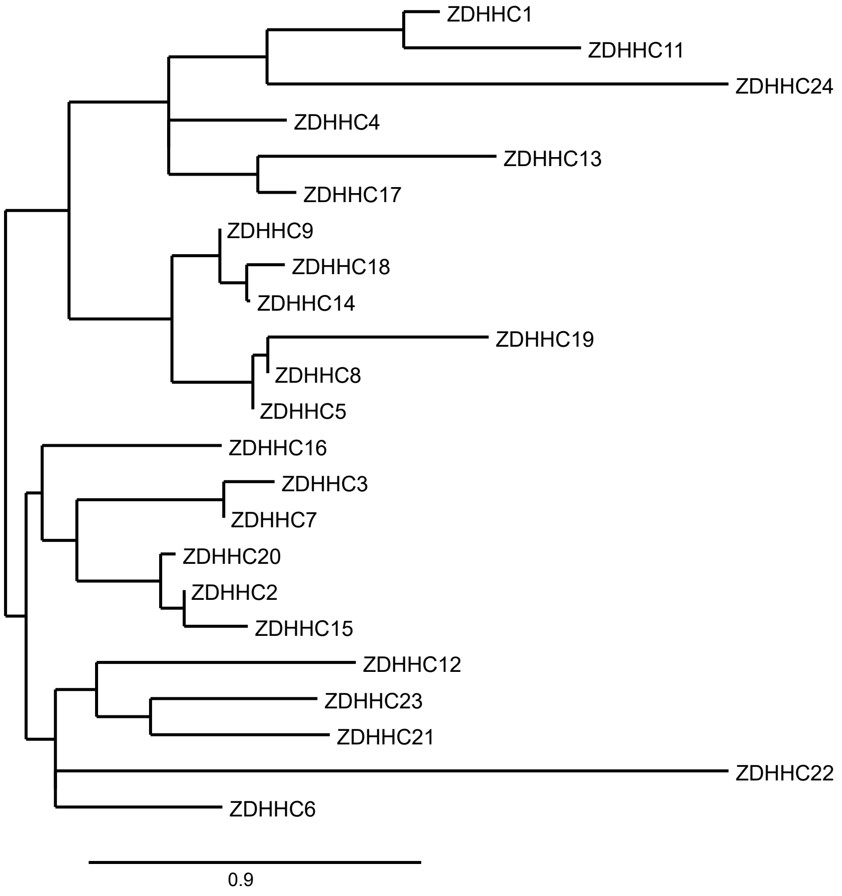


Supplementary Figure 3

***2,6-Dinitroaniline* 6, $^1$H NMR 500 MHz DMSO *$d_6$***

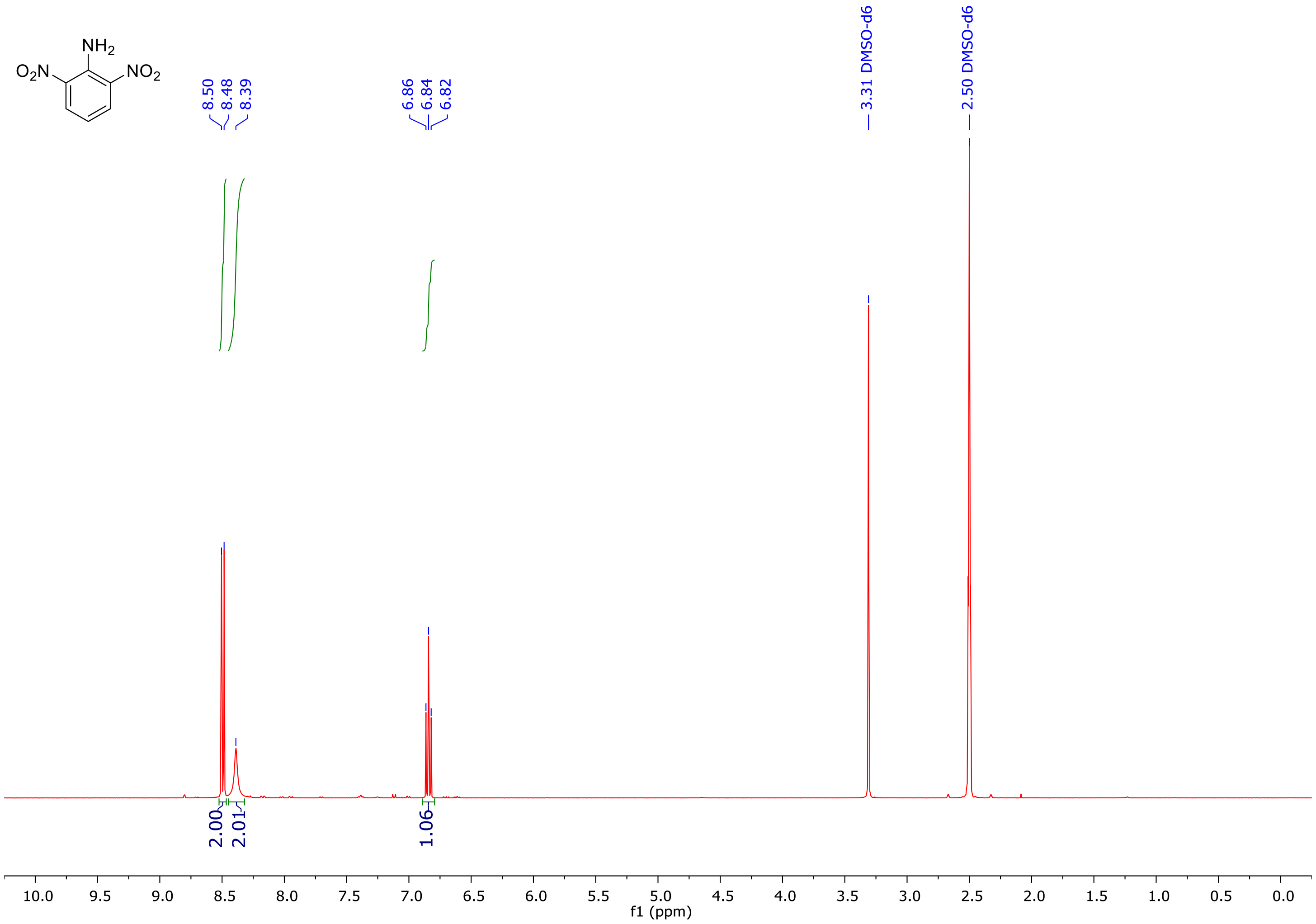

***2,6-Dinitroaniline* 6, $^{13}$C NMR 126 MHz DMSO *d*$_6$**

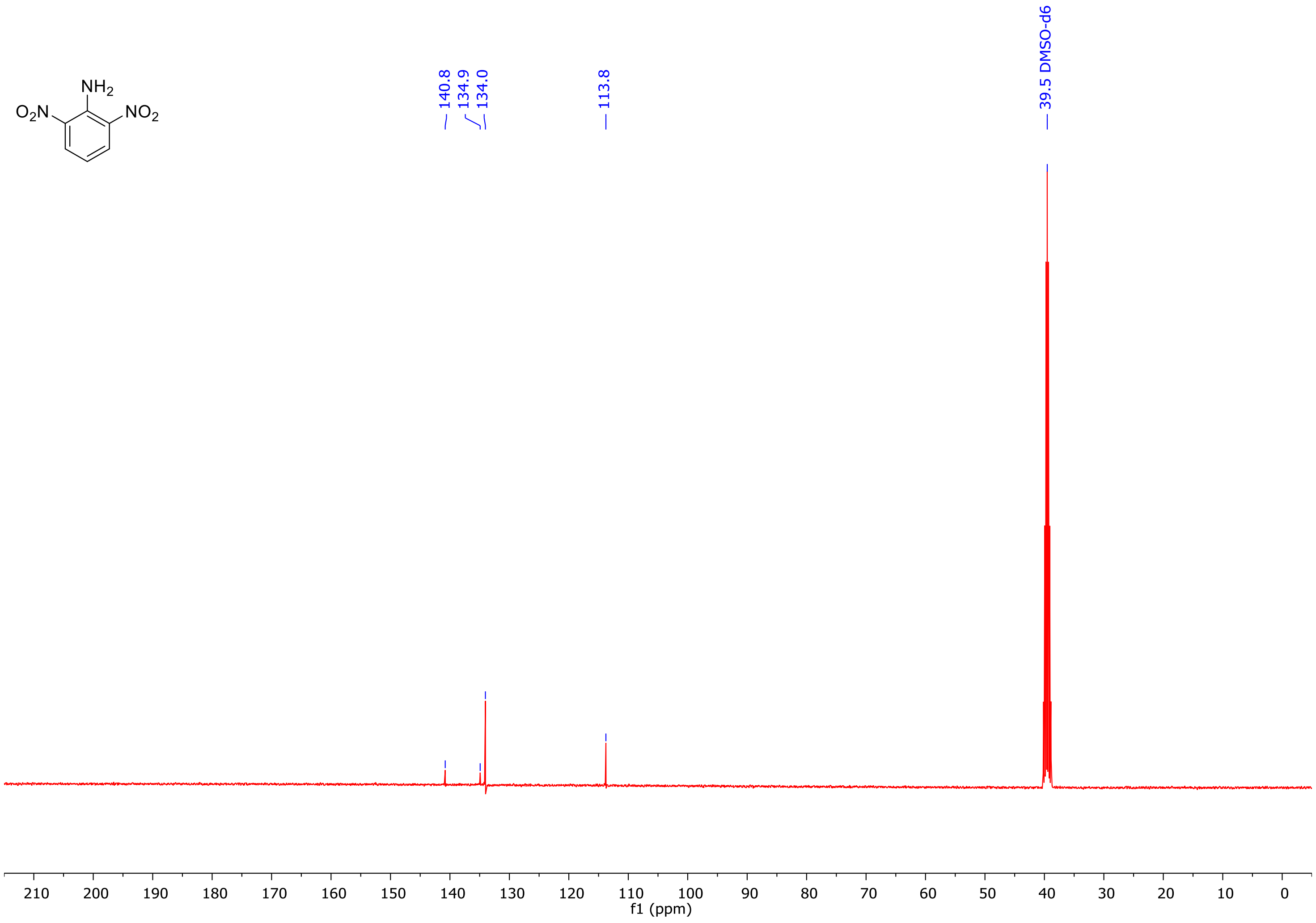

***2-Chloro-1,3-dinitrobenzene* 7, $^1$H NMR 500 MHz Acetone *d*$_6$**

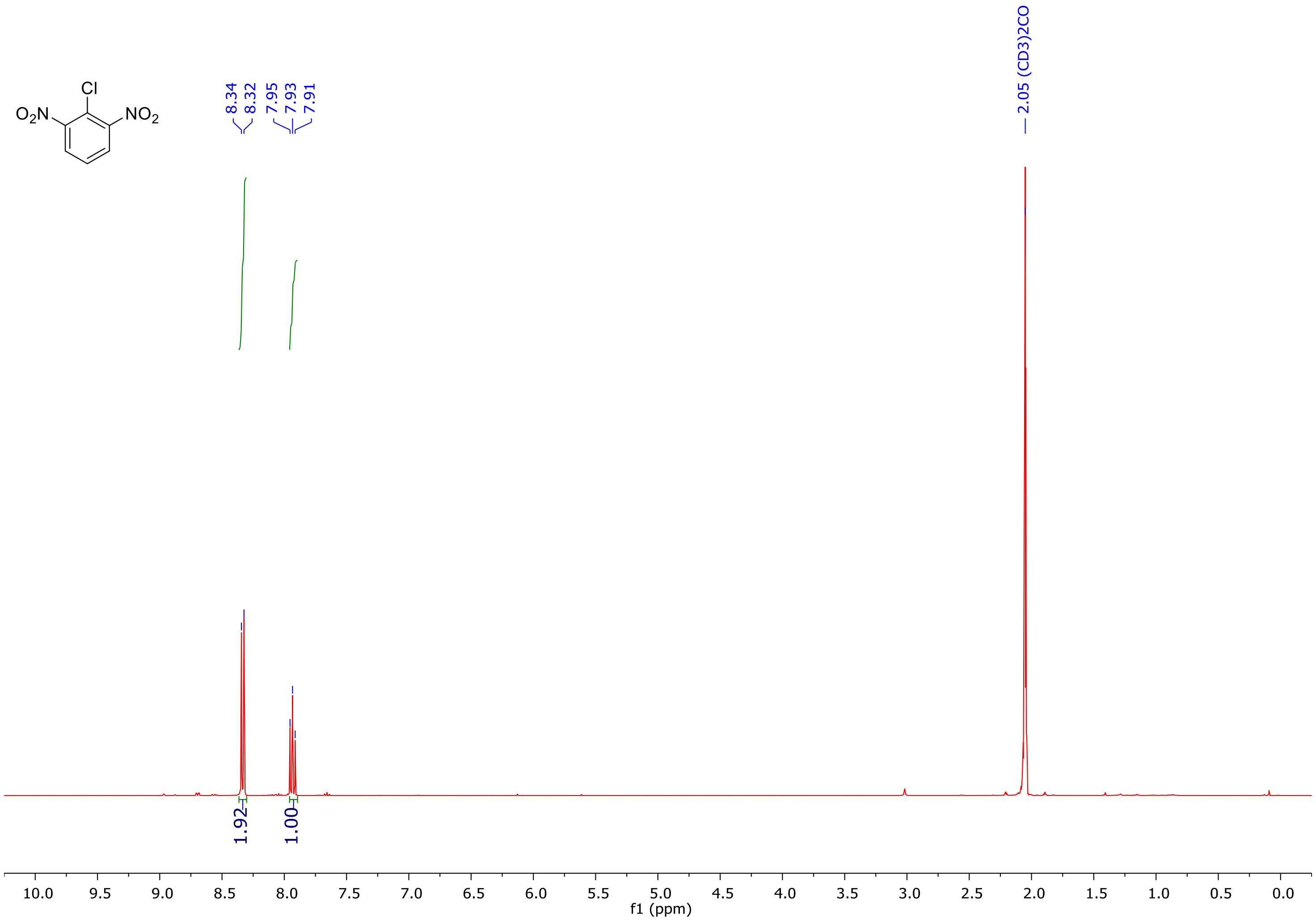

***2-Chloro-1,3-dinitrobenzene*** **7, $^{13}$C NMR 126 MHz Acetone $d_6$**

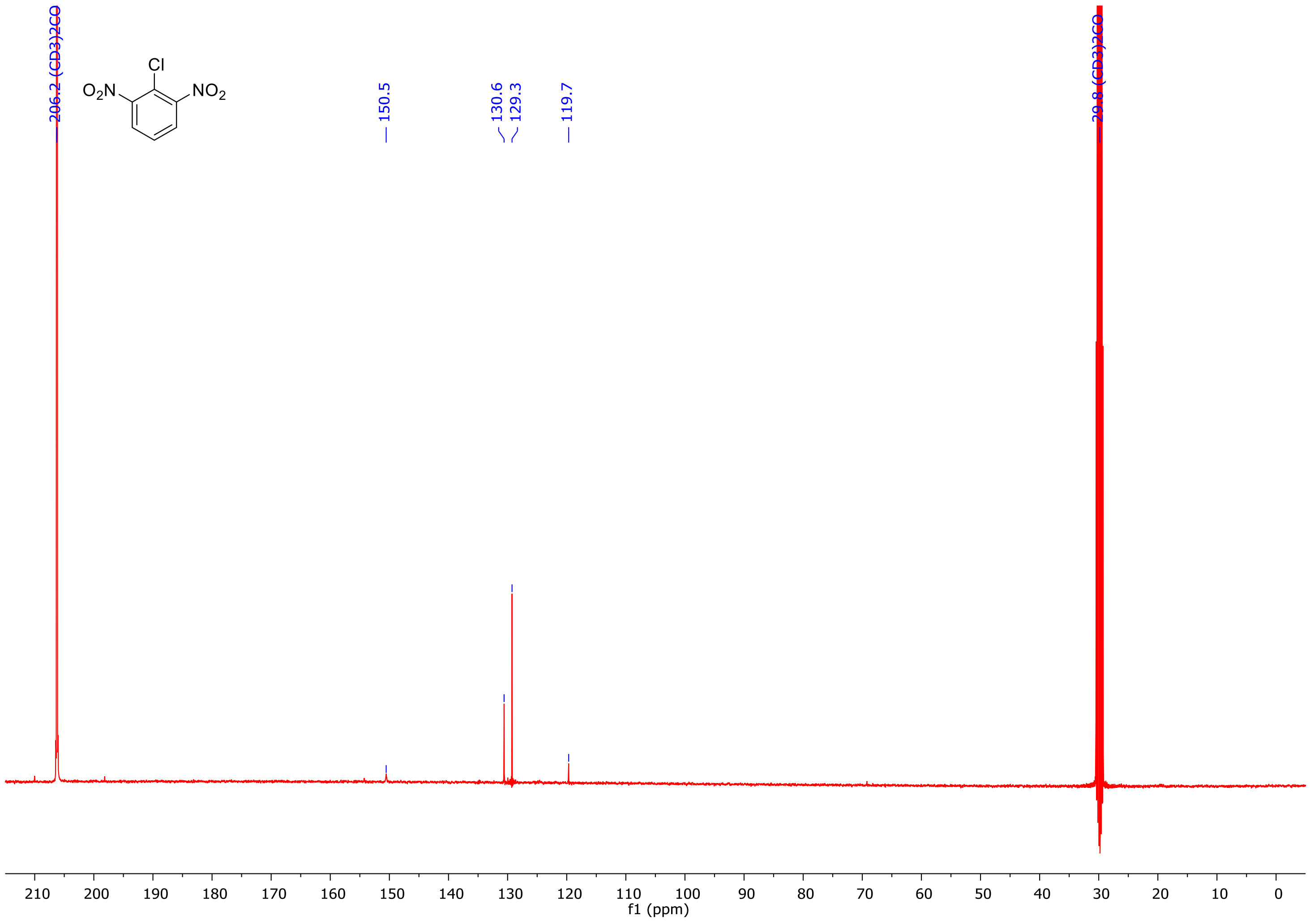

***2,6-Dinitrophenol* 8, $^1$H NMR 500 MHz Acetone *d*$_6$**

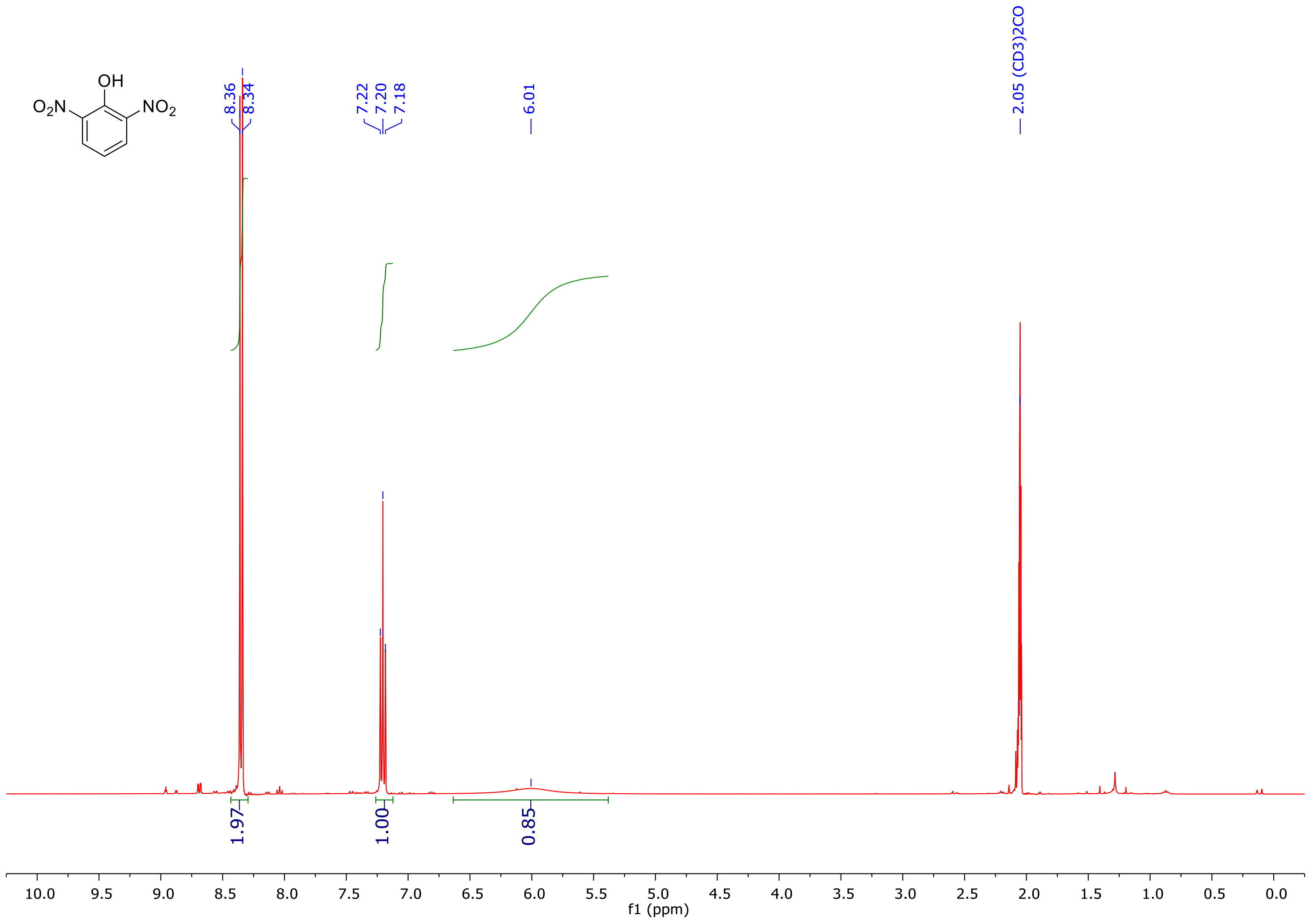

***2,6-Dinitrophenol* 8, $^{13}C$ NMR 126 MHz Acetone $d_6$**

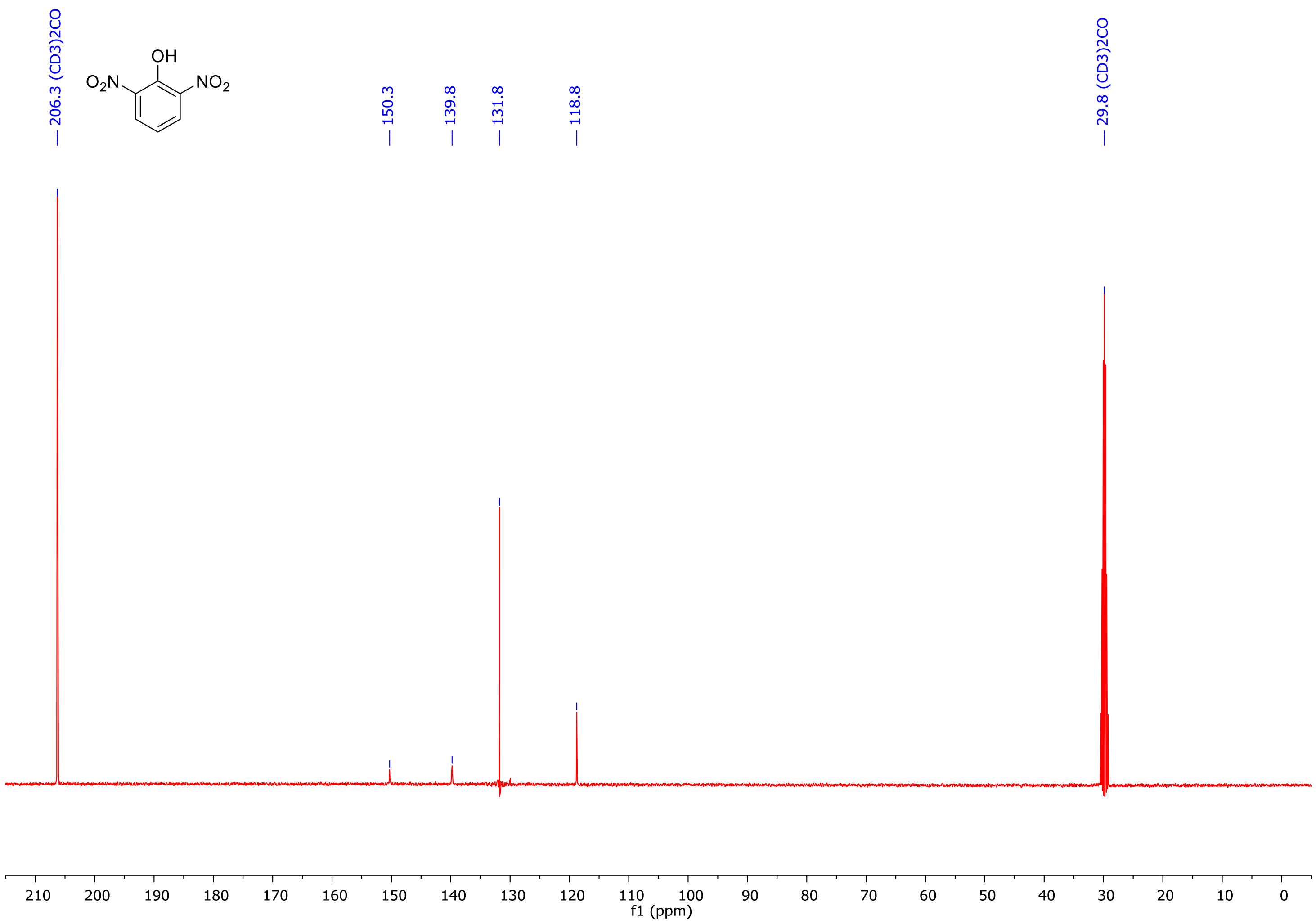

**_2-Amino-6-nitrophenol_ 9, [1]H NMR 500 MHz Chloroform _d_**

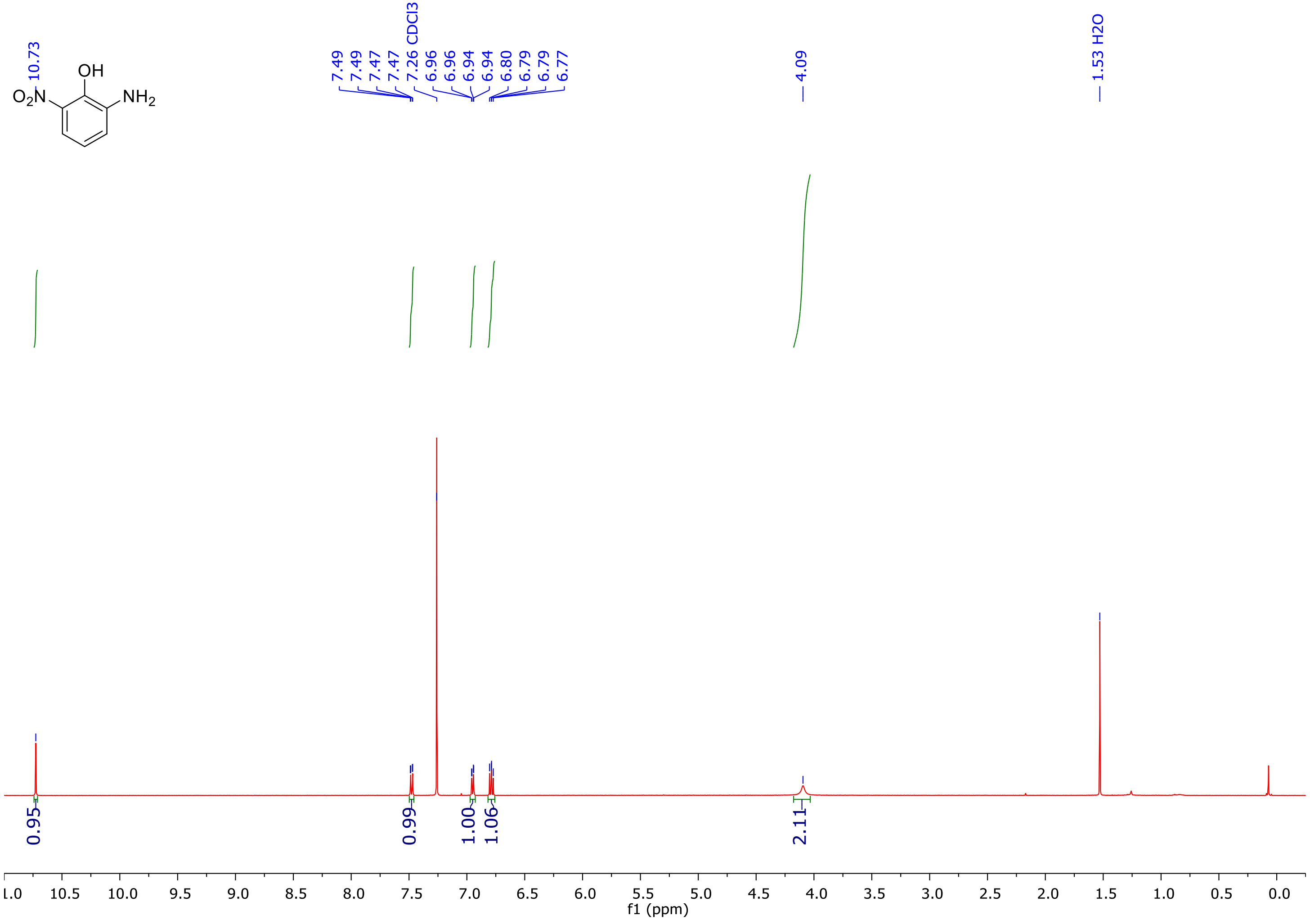

***2-Amino-6-nitrophenol* 9, $^{13}$C NMR 126 MHz Chloroform *d***

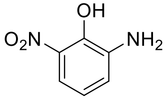

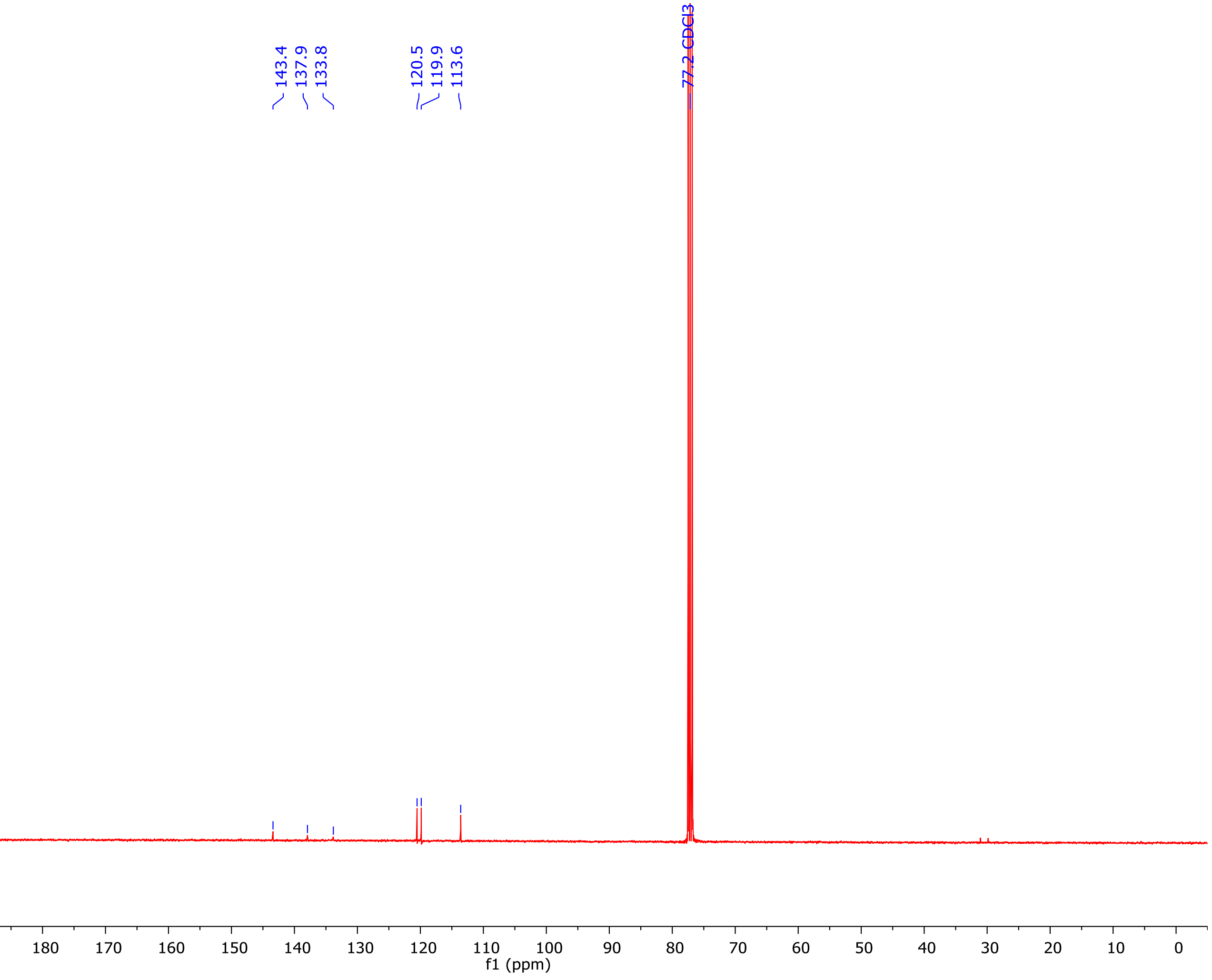

***8-Nitro-3,4-dihydro-2H-benzo[b][1,4]oxazine-2-carbonitrile*** **11, $^{1}$H NMR 500 MHz Acetone** ***$d_6$***

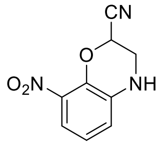


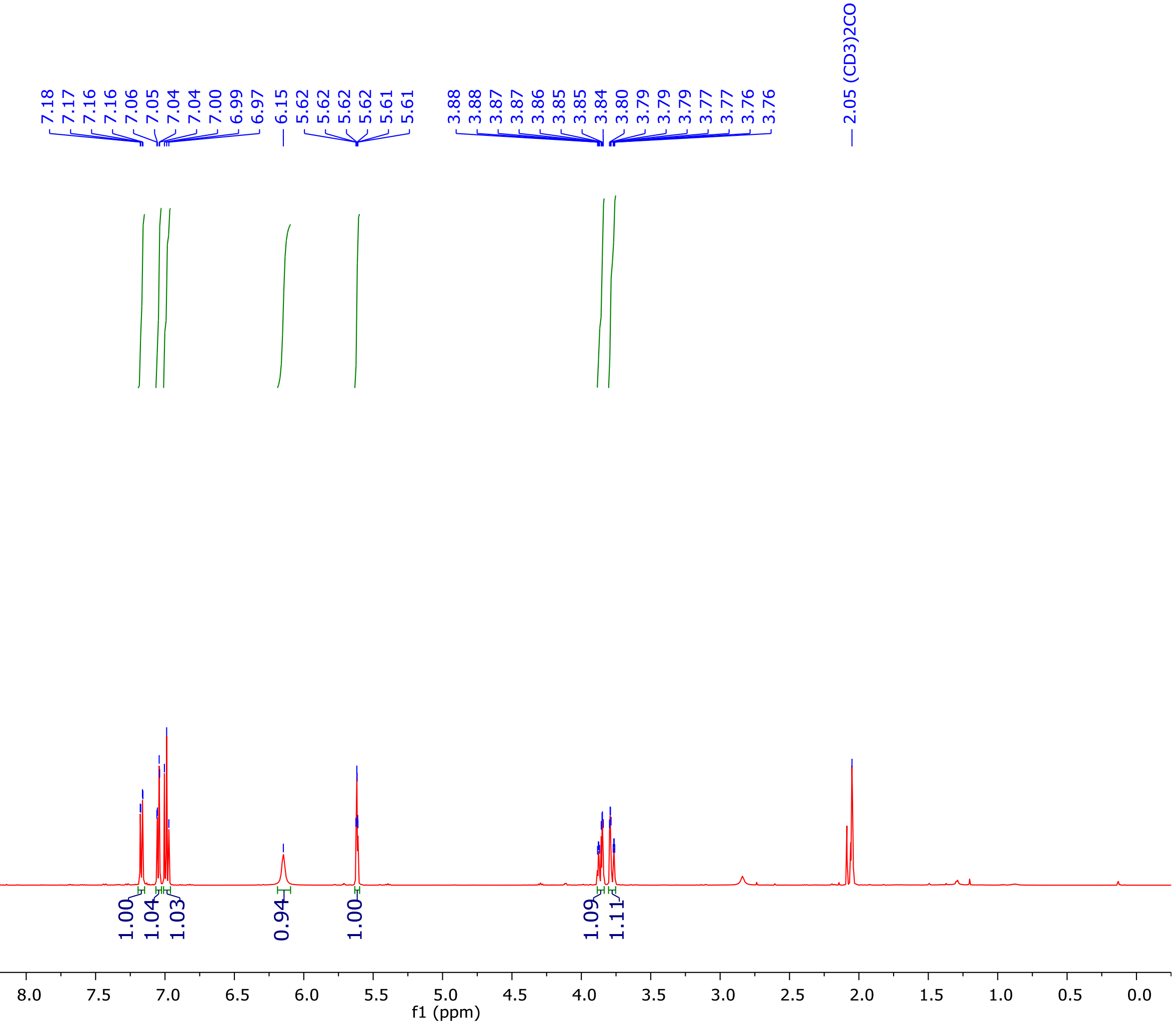

***8-Nitro-3,4-dihydro-2H-benzo[b][1,4]oxazine-2-carbonitrile* 11, $^{13}$C NMR 126 MHz Acetone *$d_6$***

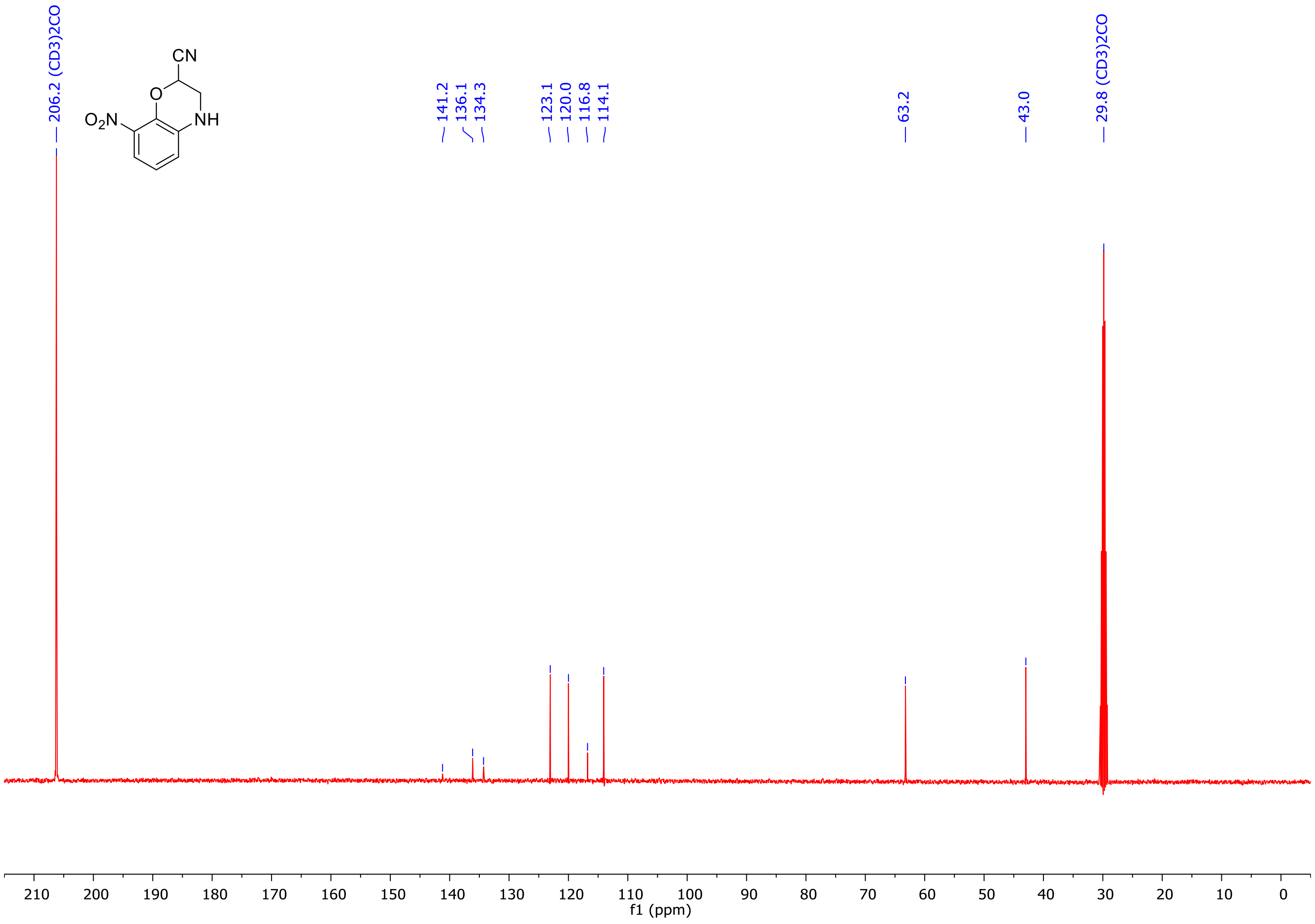

*8-Amino-3,4-dihydro-2H-benzo[b][1,4]oxazine-2-carbonitrile* 12, $^1$H NMR 500 MHz Chloroform *d*

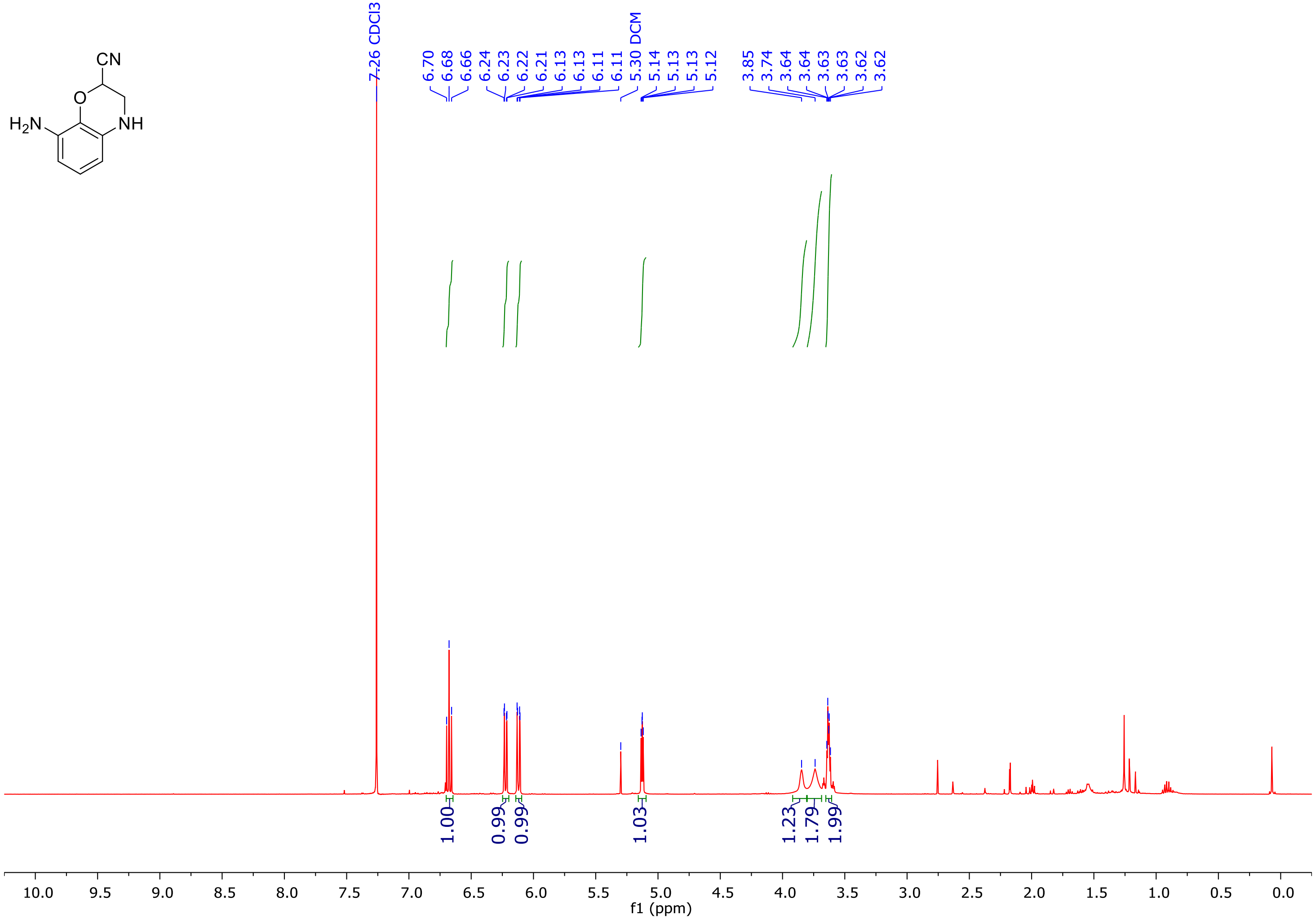

*8-Amino-3,4-dihydro-2H-benzo[b][1,4]oxazine-2-carbonitrile* 12, $^{13}$C NMR 126 MHz Chloroform *d*

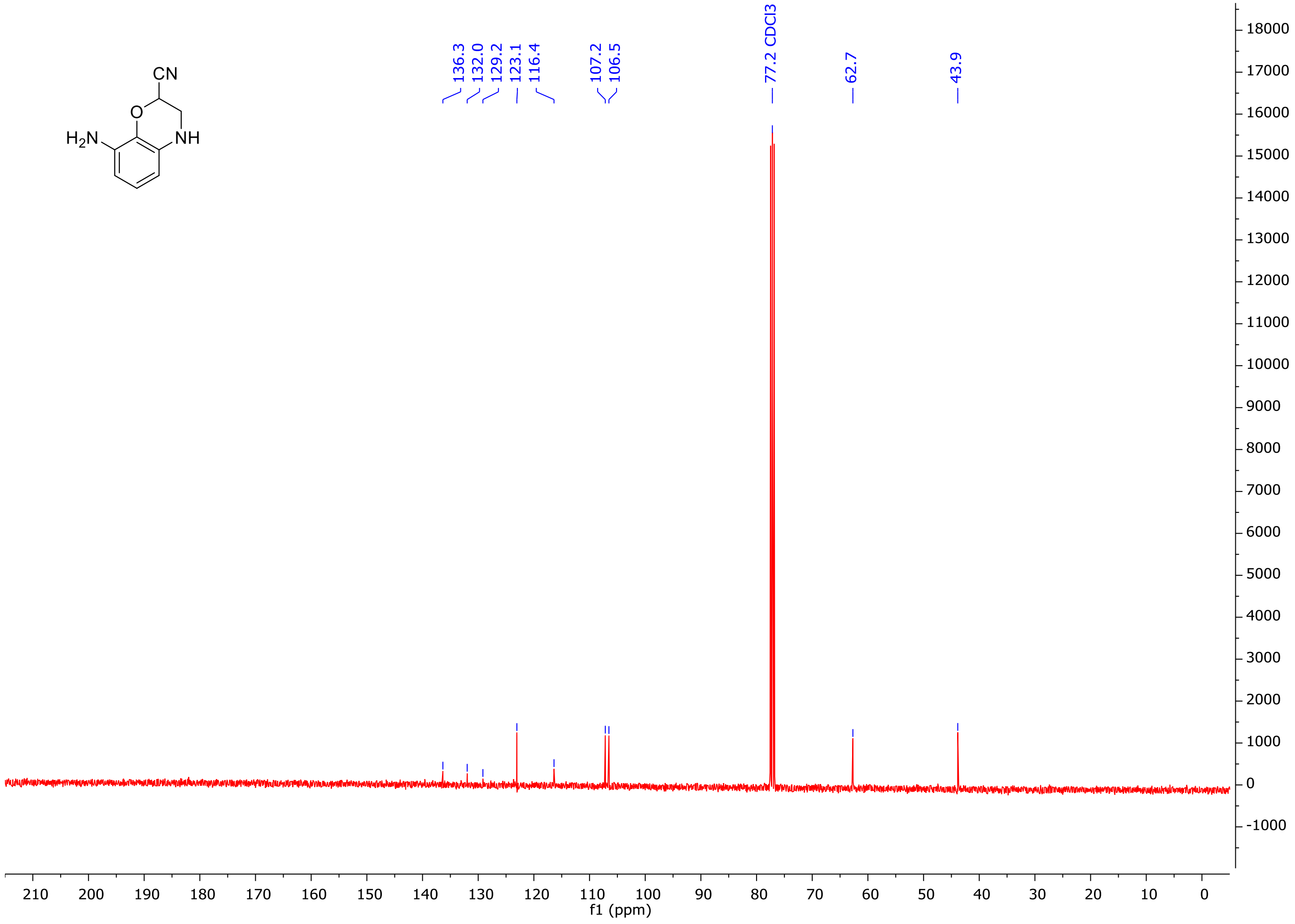

*N-(2-Cyano-3,4-dihydro-2H-benzo[b][1,4]oxazin-8-yl)-4-(heptyloxy)benzamide* 15, $^{1}$H NMR 400 MHz Acetone $d_6$

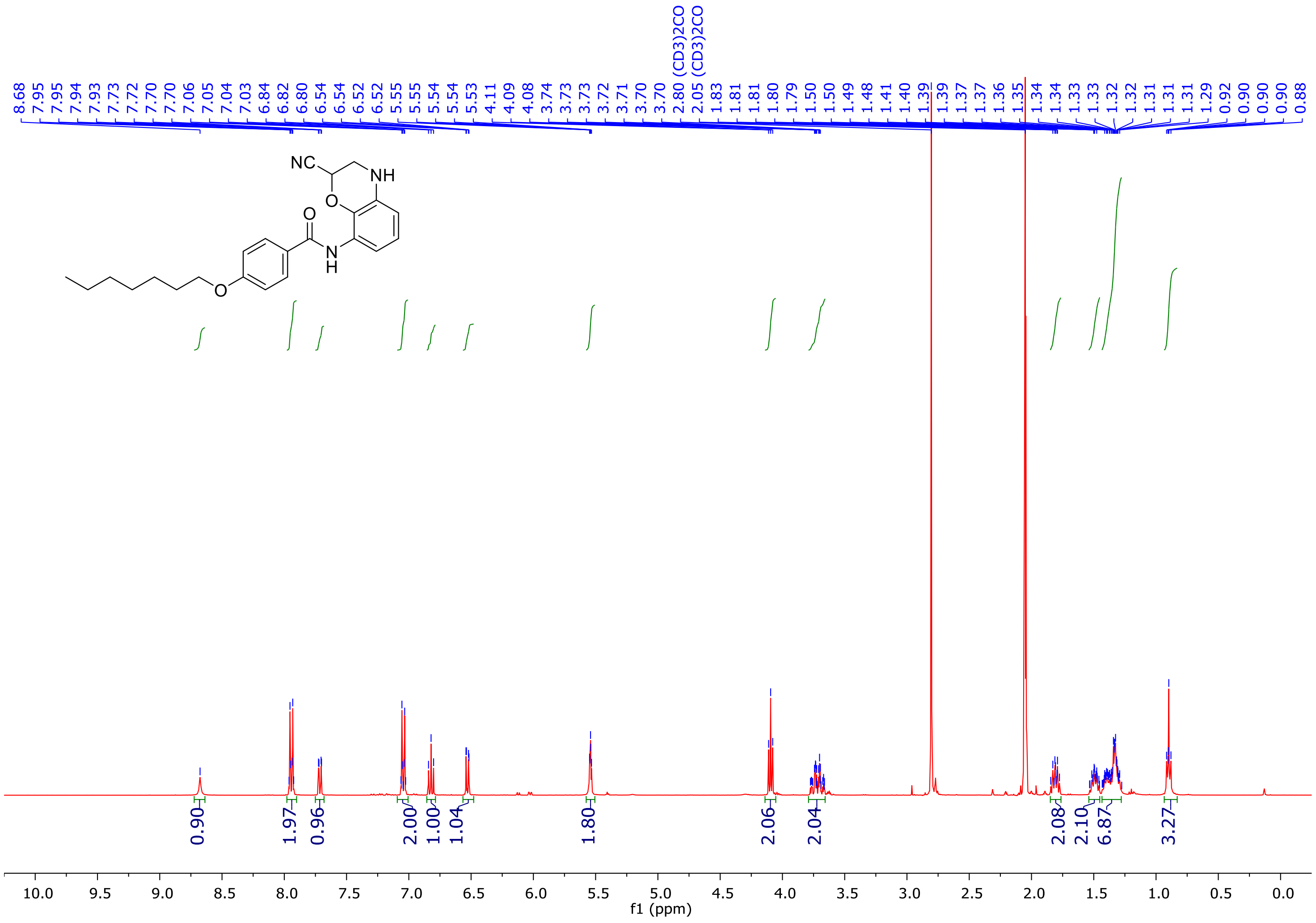

***N*-(2-Cyano-3,4-dihydro-2*H*-benzo[b][1,4]oxazin-8-yl)-4-(heptyloxy)benzamide** 15, $^{13}$C NMR 101 MHz Acetone $d_6$**

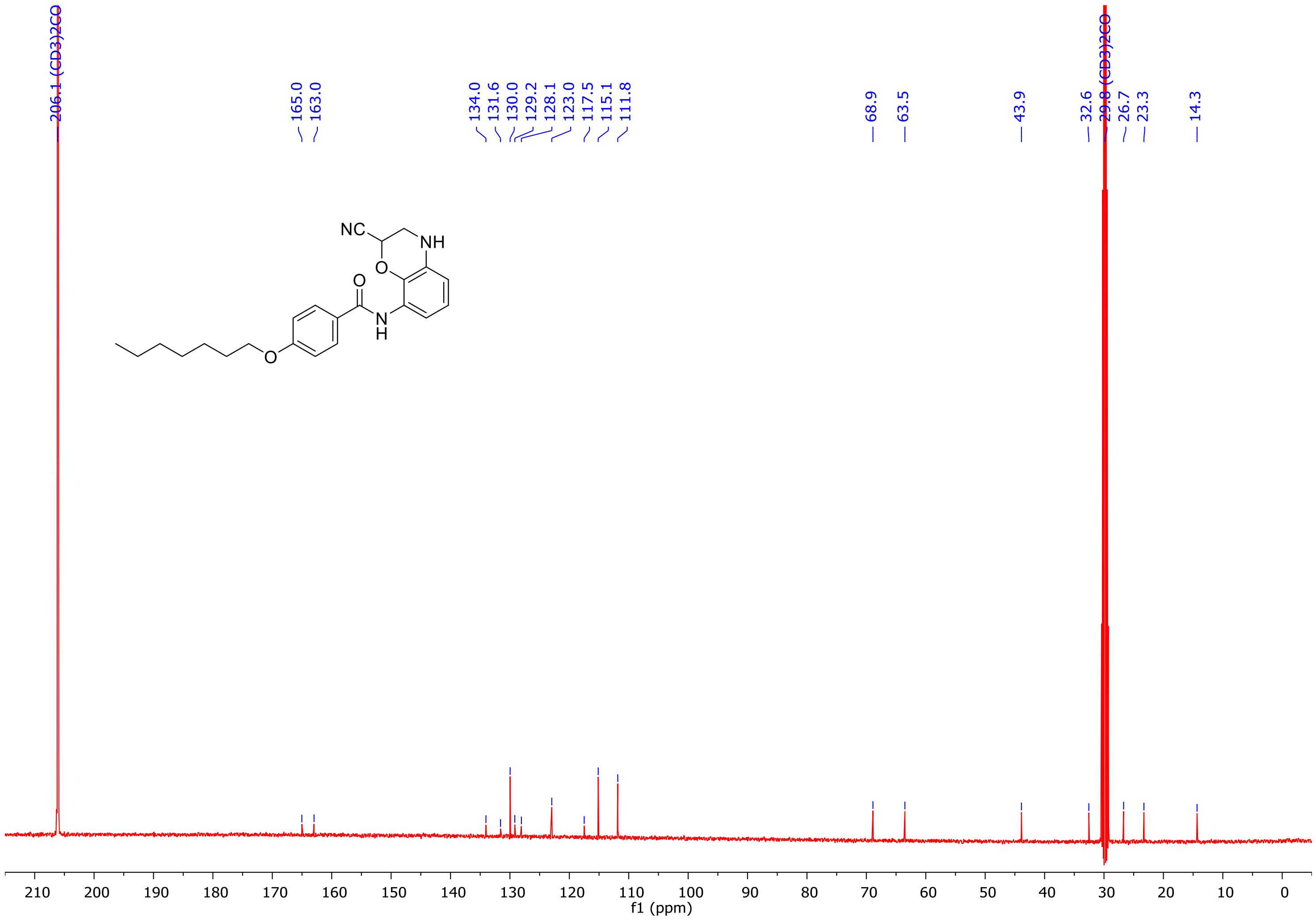

***N*-(2-Cyano-3,4-dihydro-2*H*-benzo[*b*][1,4]oxazin-8-yl)-4-(4-phenylbutoxy)benzamide** **16**, $^{1}$H NMR 400 MHz Acetone $d_6$

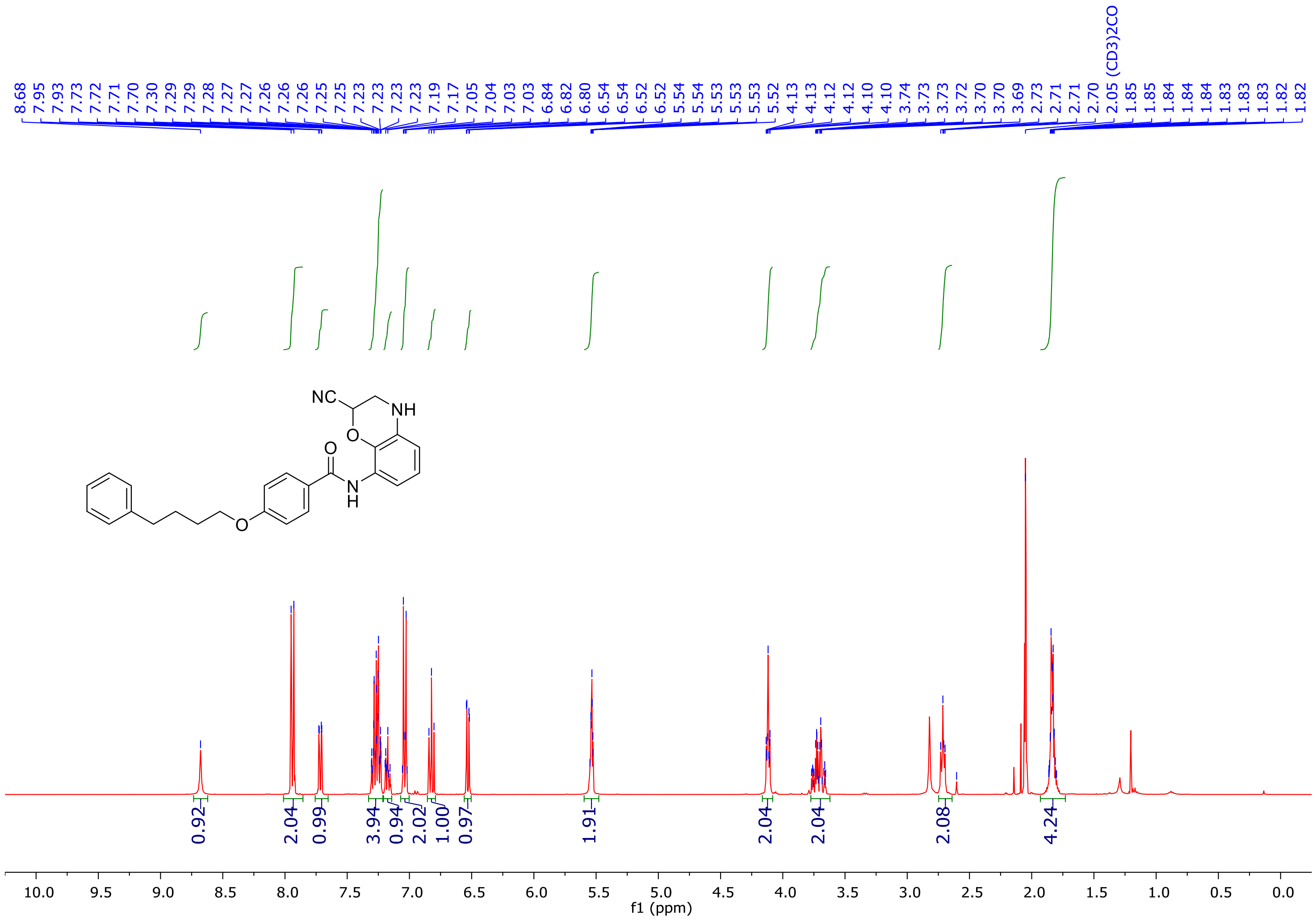

***N*-(2-Cyano-3,4-dihydro-2*H*-benzo[*b*][1,4]oxazin-8-yl)-4-(4-phenylbutoxy)benzamide** **16, $^{13}$C NMR 101 MHz Acetone *d*$_6$**

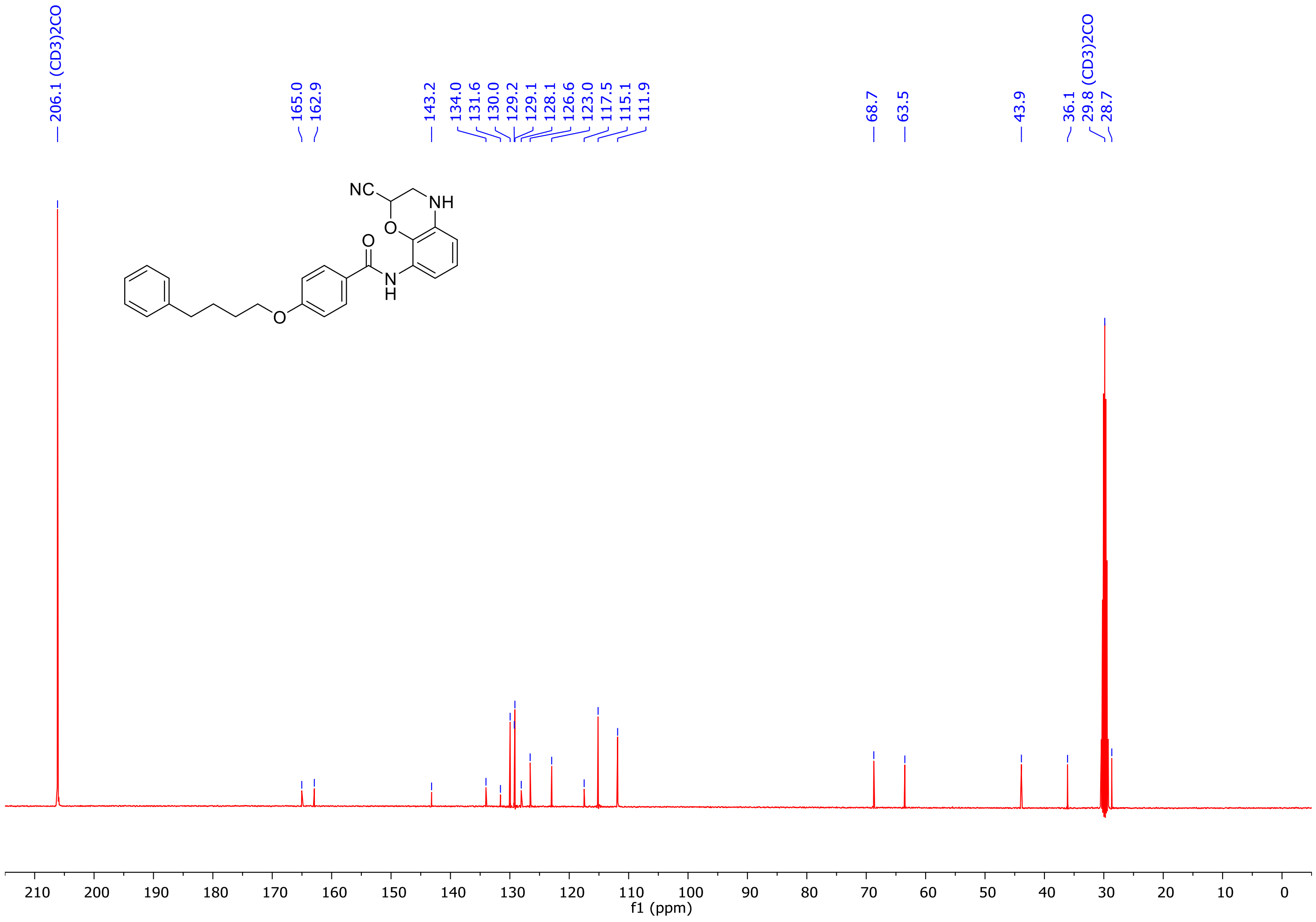

***N*-(2-(1*H*-Tetrazol-5-yl)-3,4-dihydro-2*H*-benzo[b][1,4]oxazin-8-yl)-4-(heptyloxy)benzamide** **TTZ-1 (1), $^1$H NMR 400 MHz Acetone *d*$_6$**

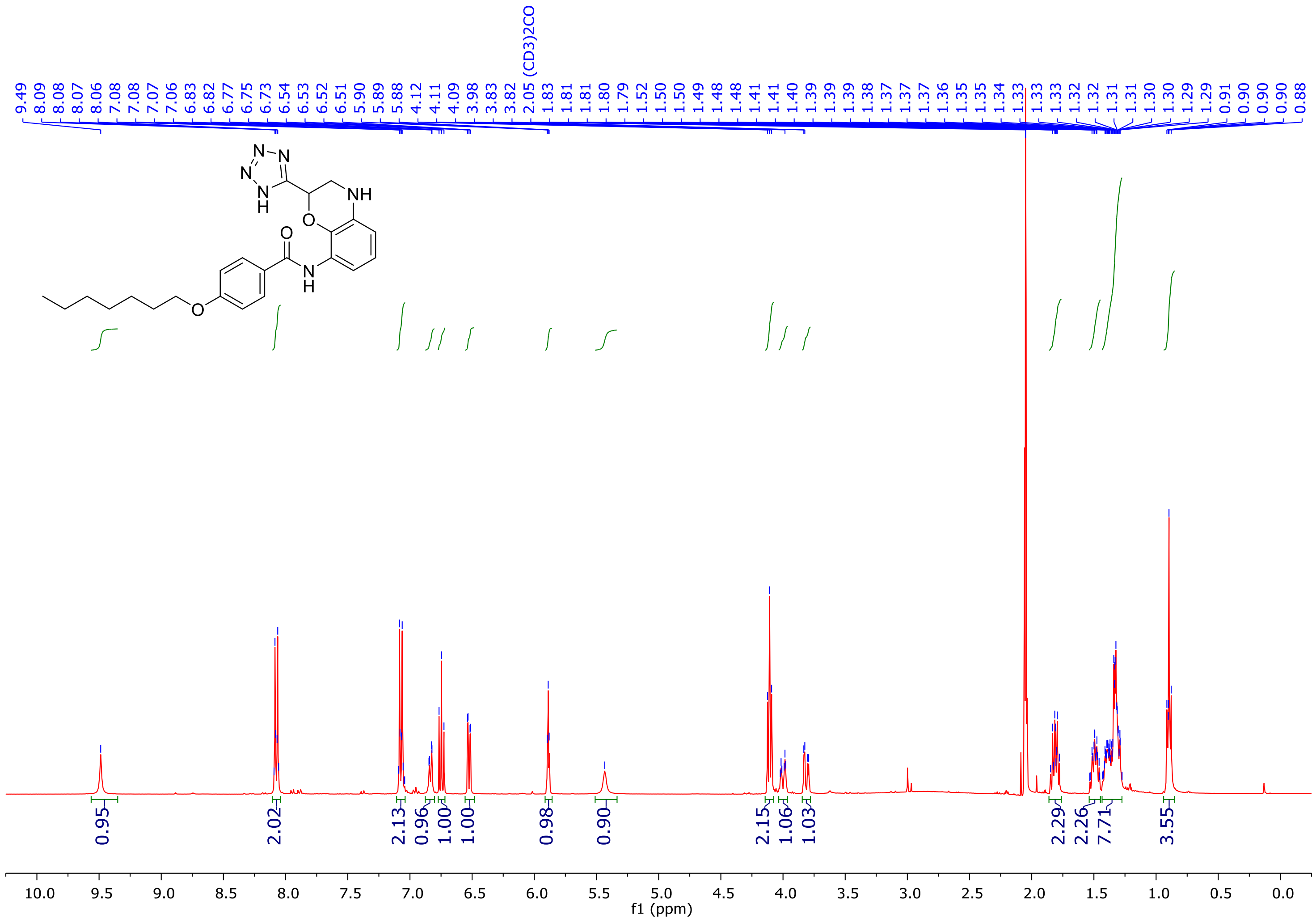

***N*-(2-(1*H*-Tetrazol-5-yl)-3,4-dihydro-2*H*-benzo[b][1,4]oxazin-8-yl)-4-(heptyloxy)benzamide** **TTZ-1 (1), $^{13}$C NMR 101 MHz Acetone $d_6$**

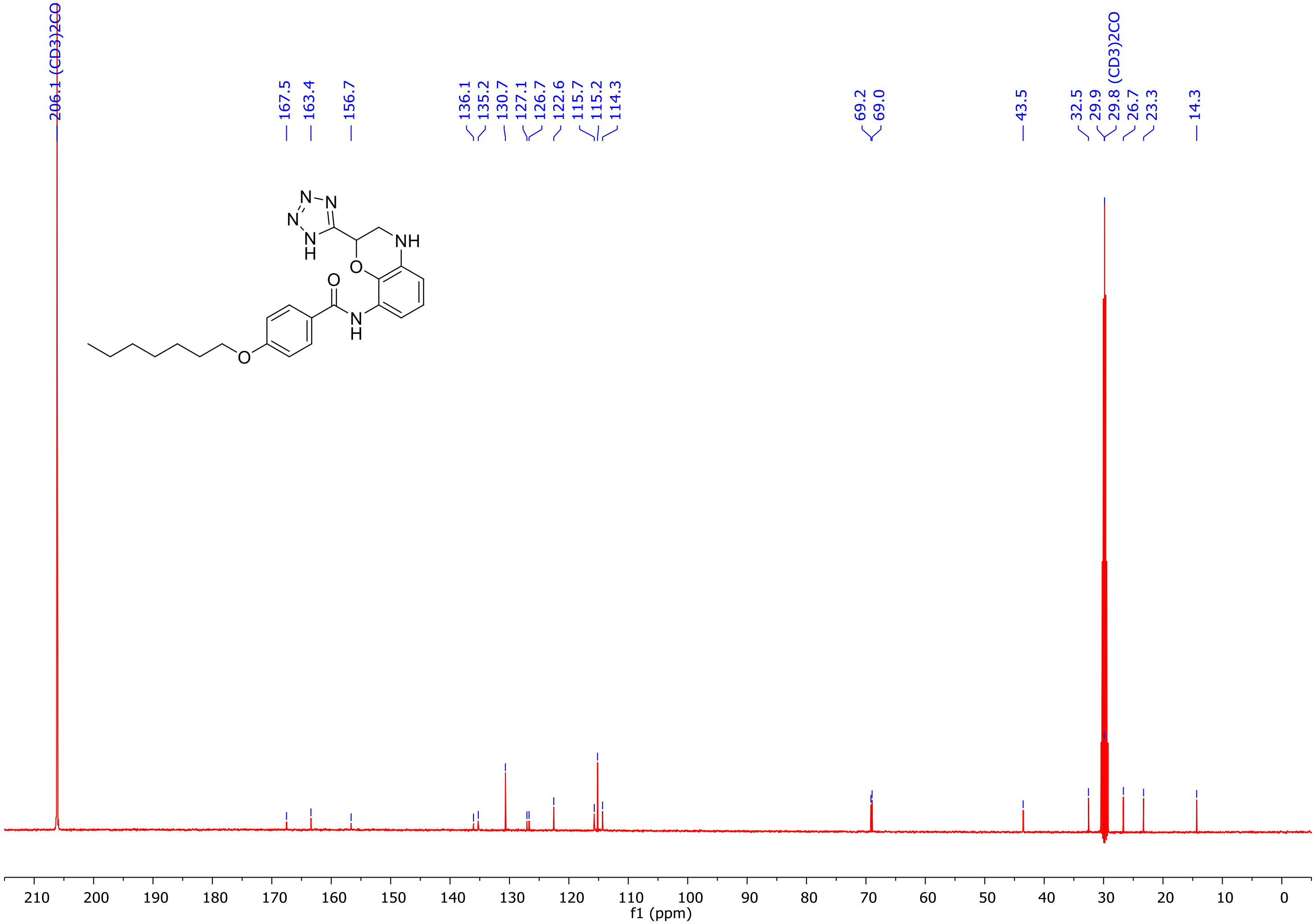

***N*-(2-(1*H*-Tetrazol-5-yl)-3,4-dihydro-2*H*-benzo[b][1,4]oxazin-8-yl)-4-(4-phenylbutoxy)benzamide** **TTZ-2 (2), $^1$H NMR 400 MHz Acetone $d_6$**

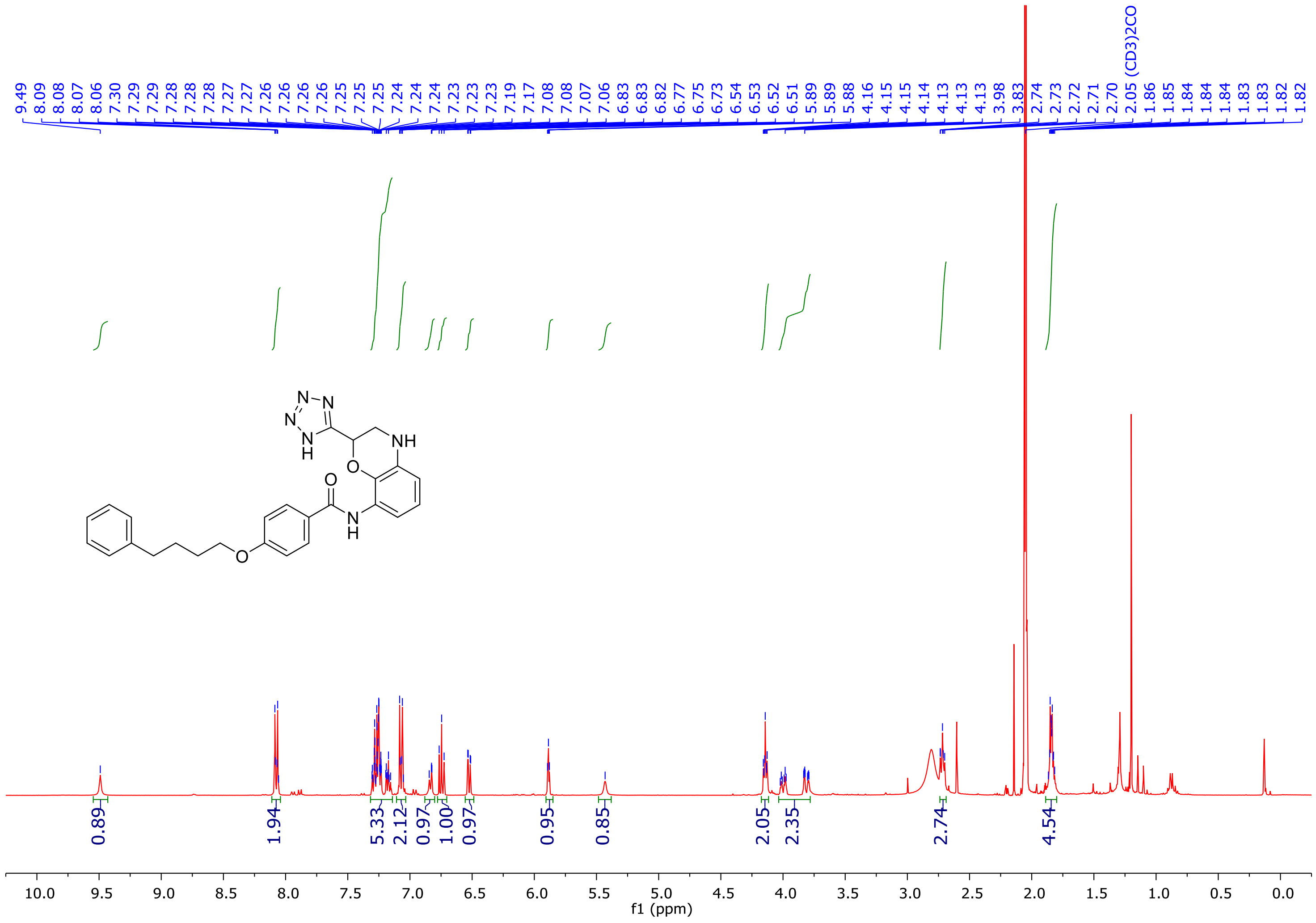

***N*-(2-(1*H*-Tetrazol-5-yl)-3,4-dihydro-2*H*-benzo[b][1,4]oxazin-8-yl)-4-(4-phenylbutoxy)benzamide** **TTZ-2 (2), $^{13}$C NMR 101 MHz Acetone *d*$_6$**

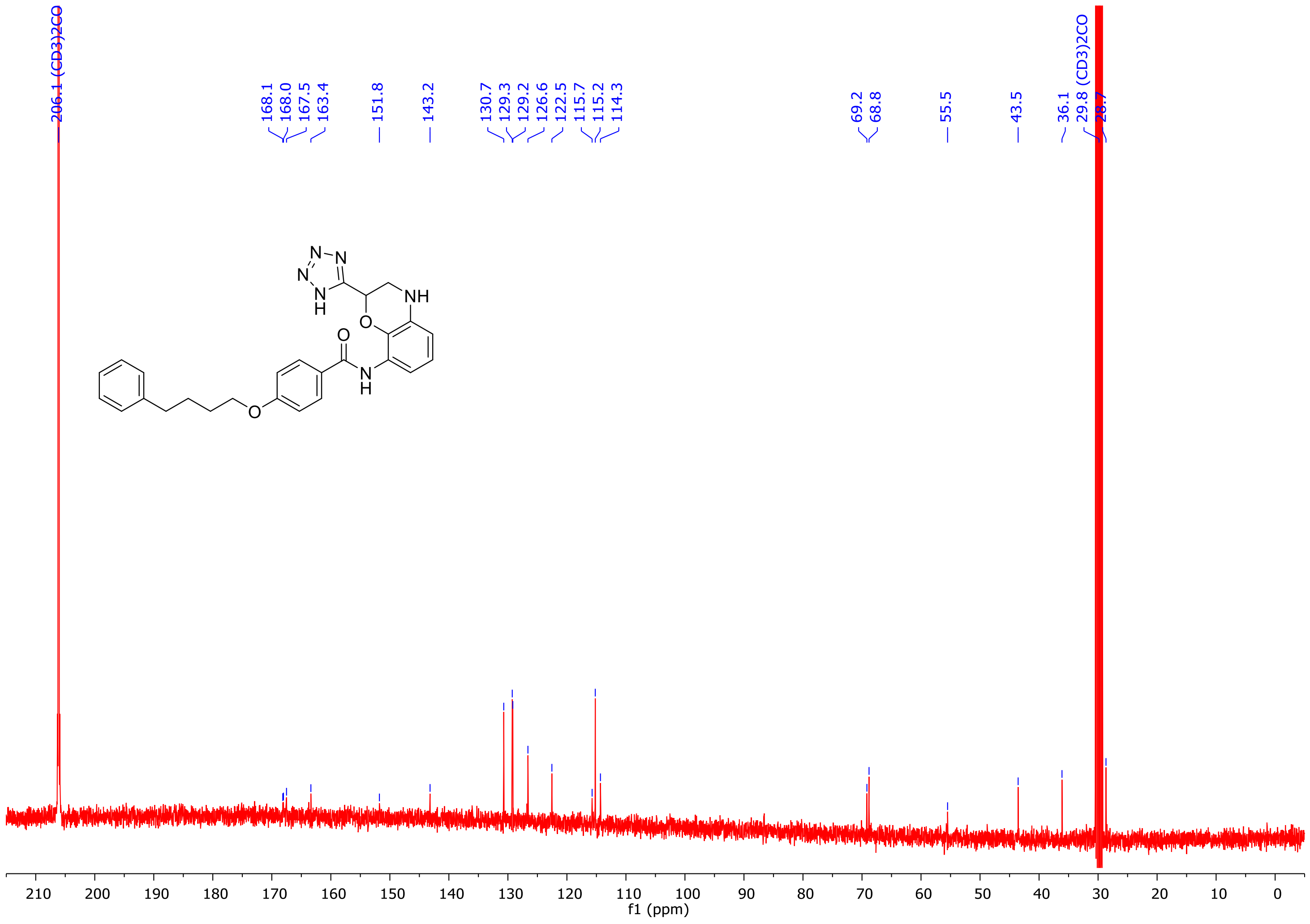
